# The DoGA Consortium Atlas of Canine Enhancers and Promoters Across Tissues and Development

**DOI:** 10.64898/2026.03.24.713649

**Authors:** Isil Takan, Matthias Hortenhuber, Noora Salokorpi, Rijja Bokhari, Cesar Araujo, Rasha Aljelaify, Ileana Quintero, Sini Ezer, Faezeh Mottaghitalab, Amitha Raman, Fiona Ross, DoGA Consortium, Tarja Jokinen, Pernilla Syrja, Danika Bannasch, Antti Iivanainen, Marjo K Hytonen, Juha Kere, Hannes Lohi, Carsten O. Daub

## Abstract

The domestic dog is a powerful genetic model for complex traits, disease and behaviours relevant to human biology, yet the canine genome still lacks a systematic transcription-defined atlas of regulatory elements. Existing resources have improved annotation but largely infer regulatory activity from chromatin features rather than directly mapping transcription initiation at promoters and enhancers. Here, we address this gap using 114 CAGE-seq libraries spanning 56 tissues and developmental stages from 9 dogs. We identify 68,446 promoters and 46,661 active enhancers, and define their tissue-enriched activity across the canine body. Regulatory programmes are organized around shared transcription factor motif infrastructures, with prominent enrichment of KLF family motifs across tissues. The cerebellum emerges as a major regulatory hub, showing the highest density of enhancer-promoter interactions among brain regions. During embryogenesis, enhancer activity shifts from early neurodevelopmental patterning programmes, including OTX2 and ZIC1/4, towards regulators of functional maturation and synaptic organization, including AQP4 and NLGN3. Comparative analyses revealed 69 enhancers with highly similar regulatory structures linked to potential orthologous genes in dog and human. Together, these data establish a tissue-resolved transcription-based atlas of the canine regulatory genome that advances functional annotation and improves interpretation of non-coding variation in comparative and disease genomics.

## Introduction

The domestic dog (*Canis lupus familiaris*) has emerged as a powerful genetic model for dissecting complex traits, diseases, and behavior relevant to human biology. Strong artificial selection during recent breeding history has produced long linkage disequilibrium blocks within breeds and pronounced genetic structure between breeds, enabling efficient mapping of heritable phenotypes while maintaining direct translational relevance to humans^1–4^.

Improvements in canine reference genomes, from the original Boxer assembly to CanFam3.1 and the long-read–based CanFam4 reference, have substantially refined gene models and transcript annotation^5–7^. However, these annotations rely largely on conventional RNA sequencing, which is not designed to resolve transcription start sites (TSSs) at base-pair resolution or to directly identify active distal regulatory elements. As a result, promoter architecture, alternative promoter usage, and enhancer landscapes remain incompletely defined in the dog genome.

Recent functional genomics resources have begun to address this limitation. BarkBase provides multi-tissue RNA-seq, whole-genome sequencing, and chromatin accessibility profiles^8^, while the EpiC Dog project integrates transcriptomes with histone modifications and DNA methylation to derive chromatin-state maps and inferred regulatory elements^9^. These efforts represent major advances for canine epigenomics but primarily infer regulatory activity indirectly from chromatin features rather than from transcription initiation itself.

In contrast, large-scale human and mouse studies have shown that transcription initiation–based annotation is essential for comprehensive regulatory maps. Using Cap Analysis of Gene Expression (CAGE), the FANTOM5 project identified promoters at single-nucleotide resolution and discovered active enhancers through bidirectional transcription across hundreds of tissues and cell types^10^. These data demonstrated strong tissue specificity of regulatory activity and revealed that disease- and trait-associated variants are highly enriched in regulatory regions ^11,12^.

Despite rapidly expanding catalogs of canine genetic variation from population-scale sequencing efforts such as Dog10K^13–15^, the dog genome lacks a systematic, transcription-defined atlas of promoters and enhancers. Current resources predict regulatory regions based on chromatin signatures, but do not establish where transcription initiates at promoters and enhancers across tissues. This gap limits functional interpretation of non-coding variants and constrains the use of dogs as models for regulatory mechanisms underlying complex traits and disease.

To address this gap, we established the Dog Genome Annotation (DoGA) consortium to generate a transcription-based functional map of the dog genome. In our previous work, we used 5′-end–targeted transcriptomics to construct a promoter and gene expression atlas across 100 canine tissues, identifying over 59,000 robust promoters, including more than 14,000 previously unannotated TSSs^16^. Here, we extend this framework by generating multi-tissue CAGE data to systematically annotate transcription start sites and active enhancer regions. By directly measuring transcription initiation, we define promoters at base-pair resolution and identify enhancers through characteristic bidirectional transcription, providing a tissue-resolved enhancerome that substantially improves functional interpretation of non-coding variation in the dog genome.

## Results

The DoGA Consortium tissue biobank includes tissue samples from 46 animals, including 13 dogs, twelve dog embryos, and 21 wolves, including eight adult wolves and 13 wolf pups^16^.

To identify active promoters and enhancers in dogs, we used 162 tissue samples from nine adult dogs and twelve dog embryos, covering 13 organ systems (Figure 1). The majority of the samples (57 out of 162 samples) were collected from the central nervous system to allow addressing brain disease and behavior-focused research.

**Figure 1:**
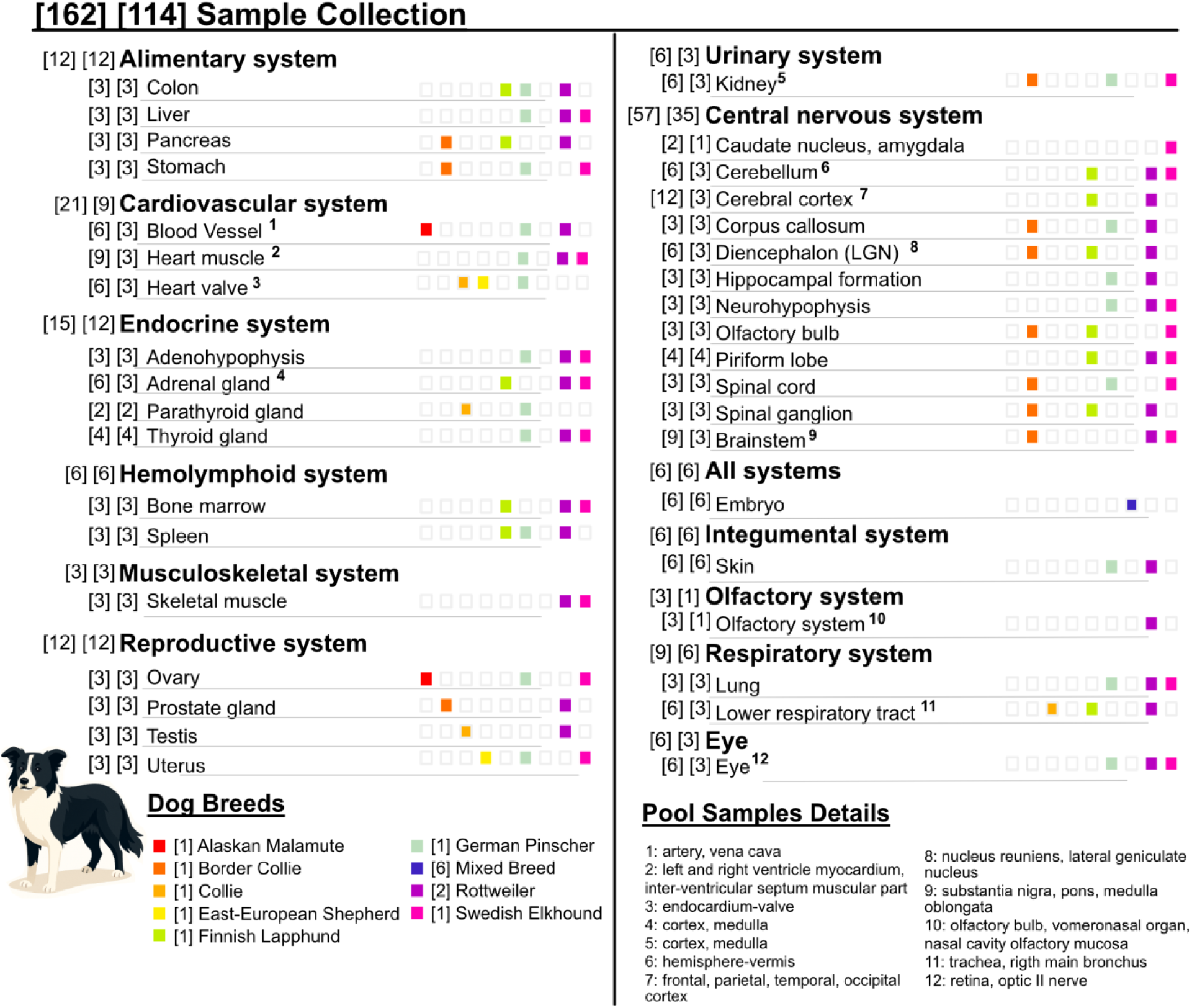
DoGA CAGE Sample Collection. The DoGA Consortium made CAGE-seq data from 114 dog samples and sample pools from nine dog breeds and twelve dog embryos. Organ systems are displayed in bold. Dog breeds are displayed in different colors. The number of tissues per CAGE sample is displayed in brackets.

We generated 114 CAGE-seq libraries from 79 individual tissue samples and from 35 pools of 83 tissue samples (on average 2 tissue samples per pool) in cases where the amount of material from a tissue was not sufficient for 1.5 micrograms of total RNA CAGE (Table S1).

### Identification of gene promoter and enhancer regions in the dog genome

The well-established CAGE sequencing method detects genome-wide mRNA starting sites in a highly sensitive and accurate way. CAGE was first published in 2003^17^ and since then served in many studies including the FANTOM^18^ and ENCODE consortia^19^ to measure the expression of transcripts and corresponding genes together with exact transcript starting sites (TSS). With the discovery that gene enhancers express enhancer RNA (eRNA)^20^, CAGE data was used to map active enhancers across human cell types and tissues^10^ and in transitioning mammalian cells^21^.

We applied the CAGE technology to 162 DoGA consortium dog samples to identify a comprehensive set of 68,446 TSS, which we refer to here as gene promoters. CAGE identified promoters in a data-driven manner without using gene or transcript models. In a subsequent step using the RefSeq set of 20,037 genes (NCBI RefSeq assembly GCF_011100685.1)^22^ as reference of the currently annotated genes in the dog genome, 53,162 of the suggested promoters corresponded to RefSeq genes. From this comprehensive promoter set, we further selected a highly expressed robust subset of 27,341 promoters consistently expressed in our DoGA samples, corresponding to 15,943 RefSeq genes. Among the promoters we identified in the DoGA samples, RefSeq did not provide a model for 15,285 of the comprehensive and 3,017 of the robust promoters. In fact, the NCBI Eukaryotic Genome Annotation Pipeline employs CAGE data for constructing gene models^22^. These non-annotated promoters constitute potentially novel transcripts and genes. Genes can be expressed from alternative promoters, and the comprehensive dataset encompassed 11,795 genes with more than one promoter while the robust promoter set contained 5,054 genes with two or more promoters (Table 1, Figure 2).

**Figure 2:**
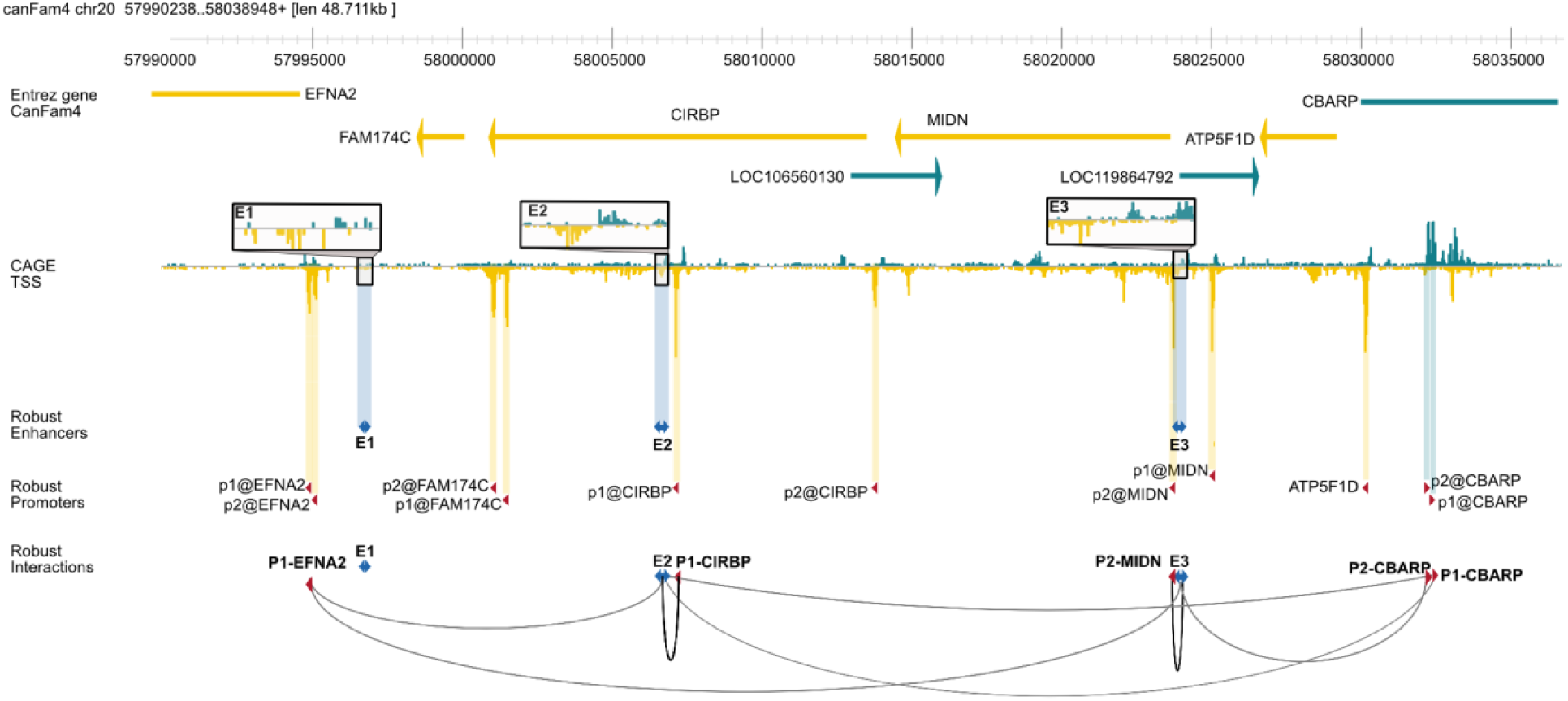
An overview of promoters and enhancers in the dog genome. CAGE transcript starting sites (TSS) together with resulting robust and permissive promoters (naming format: p1@geneA) and enhancers (naming format:E1, E2, etc.) are displayed in the context of the dog genome. Predicted regulatory interactions between enhancers and promoters are shown with arcs.

**Table 1:** DoGA promoters and enhancers in dogs from 150 dog tissue samples corresponding to 114 CAGE sequencing samples.

|  | Comprehensive | Robust |
| --- | --- | --- |
| Number of DoGA Promoters | 68,446 | 27,341 |
| Supported by ATAC | 41,894 (61%) | 19,170 (70%) |
| Supported by H3K27ac | 31,704 (46%) | 13,595 (50%) |
| Annotated to a RefSeq Gene | 53,162 (78%) | 24,324 (89%) |
| Novel DoGA Promoters<br>(compared to RefSeq genes) | 15,284 (22%) | 3,017 (11%) |
| Number of RefSeq Genes | 20,037 | 15,943 |
| With Multiple DoGA Promoters | 11,795<br>(59% of Genes) | 5,054<br>(32% of Genes) |
| With One DoGA Promoter | 8,242<br>(41% of Genes) | 10,889<br>(68% of Genes) |
| Number of DoGA Enhancers | 46,661 | 9,787 |
| Supported by ATAC | 14,805 (32%) | 4,538 (46%) |
| Supported by H3K27ac | 17,677 (38%) | 4,775 (49%) |

The DoGA CAGE dataset of 150 dog samples allowed us to further identify 46,661 comprehensive enhancer regions. In analogy to the comprehensive and robust promoters, we selected a subset of 9,787 robust enhancers displaying high expression across samples (Table 1, Figure 2).

### Promoters and enhancers are characterized by epigenetic states

Promoters and enhancers identified by CAGE are in transcriptionally active regions in the genome. These regions are characterized by open chromatin and are typically associated with epigenetic histone marks of active transcription. We compared the identified DoGA promoters and enhancers with publicly accessible data on H3K27 acetylation (H3K27Ac)^9^, the histone mark typically associated with active transcription^9,23^. Chromatin accessibility was assessed with ATAC-Seq data^8^. 13,595 (50%) of the robust DoGA promoters and 31,70 (46%) of the comprehensive DoGA promoters were supported by the active transcription mark H3K27ac (Table 1, Figure 3, Figure S10). Open chromatin ATAC-seq marks supported 41,894 (61%) of the comprehensive DoGA promoters and 19,170 (70%) of the robust DoGA promoters.

**Figure 3:**
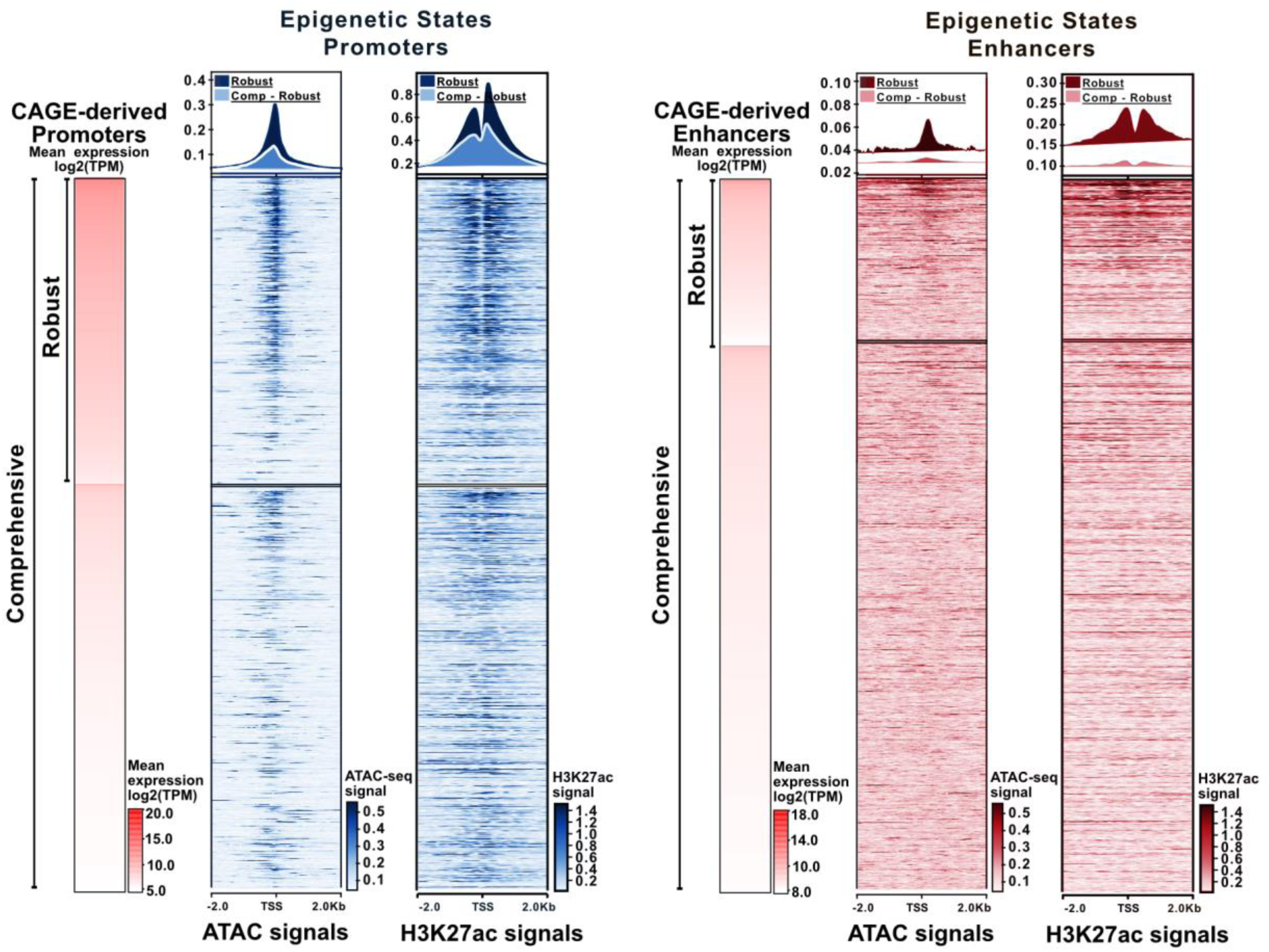
Evaluation of the DoGA enhancer and promoter candidates by open chromatin marks and histone marks. Chromatin openness ATAC-seq data from eight dog tissue samples from spleen and active transcription mark H3K27ac ChIP-seq data from eleven dog spleen samples were overlaid with DoGA promoters and enhancers (Figures S9 and S10), sorted by expression (first column).

For the DoGA enhancers, 17,667 (38%) of comprehensive and 4,775 (49%) of robust enhancers were supported by H3K27Ac. Open chromatin marks were observed for 14,805 (32%) of comprehensive and 4,538 (46%) of the robust enhancers (Table 1, Figure 3, Figure S11).

Furthermore, genomic regions around the DoGA promoters and enhancers displayed marked genome sequence conservation (Figure S15). Clear peaks were observed in ATAC-seq data, and a typical peak-valley-peak pattern appeared in H3K27ac ChIP-seq data for both promoter and enhancer candidates. The epigenetic signals were more pronounced in promoters than in enhancers.

### Promoters and enhancers cluster by tissue identity

A principal component analysis from the promoter regions (Figure 4A) and enhancer regions (Figure 4B) groups the 114 CAGE samples from 150 DoGA tissues according to organ systems and tissues. Tissue samples from the cardiovascular system were grouped together, and skeletal muscle tissue samples (part of the muscular system) and heart muscle tissues (part of the cardiovascular system) were positioned close to each other. Although located physiologically close to the central nervous system (CNS), the spinal ganglion and neurohypophysis are located further away from the central nervous system than others, while other CNS samples are grouped together. We observed similar patterns in the dimension reduction analysis based on enhancers.

**Figure 4:**
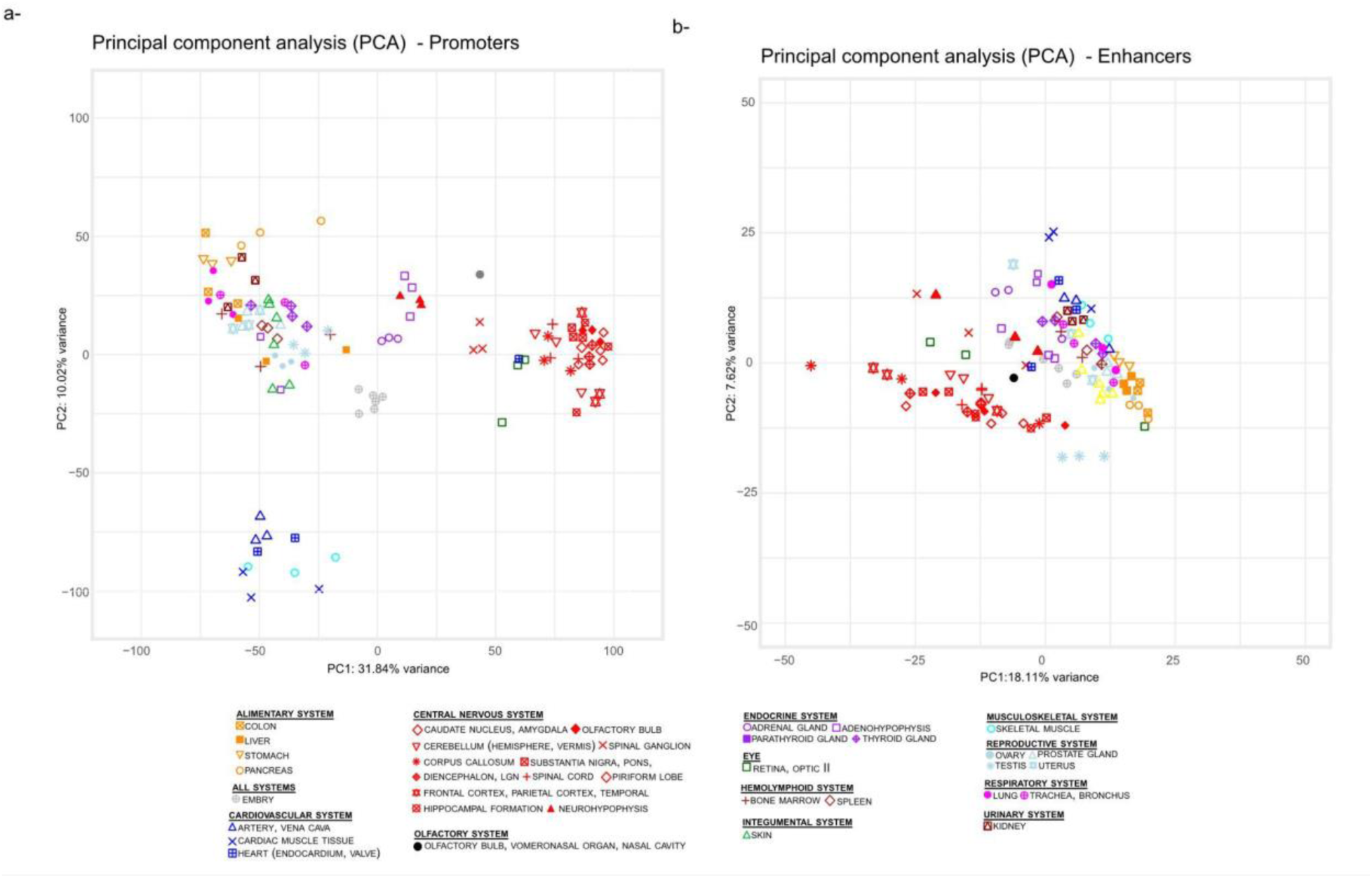
DoGA samples group according to organ systems and tissue identity. Principal component analysis (PCA) of the CAGE-seq tissue samples is based on the expression of the robust promoter set (A) or the expression of the robust enhancer set (B). Each mark indicates one sample. The color corresponds to the organ system of origin.

To determine if promoters or enhancers exhibited tissue-specific expression, we identified enriched regions where the mean expression in a target tissue exceeded the mean expression across all other tissues. To quantify this specificity, we calculated a Tissue Enrichment Score (TES) using the log-2 fold-change of expression in the target tissue relative to the average of the remaining tissues (Figures S3-S5). For instance, a TES of 3 corresponds to an 8-fold (or 2^3) increase in expression for a given tissue (Figure 5).

**Figure 5.**
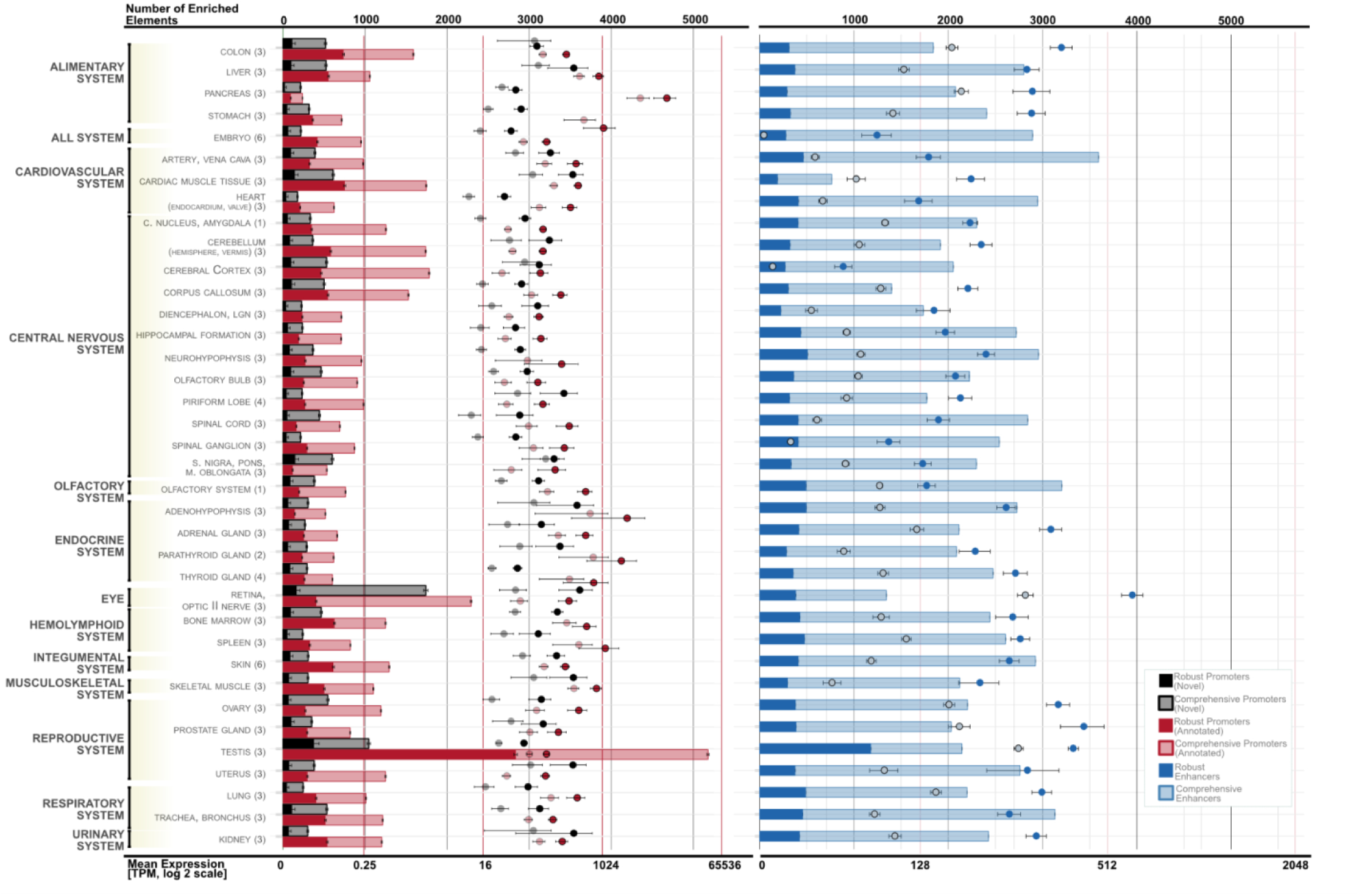
Tissue enrichment of promoter and enhancer candidates based on tissue enrichment score 3. The figure displays enhancers and promoters with tissue enrichment scores above 3 by tissue. The y-axis shows CAGE tissue pools and the related organ systems, listed alphabetically. The top x-axis indicates the number of elements enriched in each tissue, while the bottom x-axis shows the mean expression (TPM) with log 2 scale of elements enriched in those tissues.

The tissues with the most enriched promoters were the reproductive system, eye, urinary, and musculoskeletal systems; the testis had the highest number of 3,214 enriched promoters. The regions with many enriched promoters have, conversely, fewer common promoters compared to other regions (Figure S6, S7). However, the mean expression levels of tissue-enriched annotated promoters were highest in the endocrine system. The eye and testis harbored the most tissue enriched comprehensive promoters, both for annotated and novel, while the testis harbored the most tissue-enriched robust promoters.

Across all tissues, we observed a larger number of tissue-enriched enhancers compared to promoters. Similar to the promoter regions, most tissue-enriched robust enhancers were observed in the testis (1,177 enhancers) (Figure 5). The overall expression levels (normalized TPM) of enhancers show a range with less fluctuation due to their low-level expression.

Notably, the expression levels of enhancers were high in the eye, despite their relatively small number. The number of common enhancers across all tissues is relatively low in comparison to promoters.

### Enhancers regulate genes

When a promoter is under the active regulation of an enhancer, this activity can be reflected in the correlation of expression levels between the enhancer and promoter for the relevant tissue samples. In turn, potential interactions between enhancers and promoters can be detected through expression correlation analysis. The enhancer and promoter may be located at relatively long distances from each other on the same chromosome. Such proximity can be assessed with chromatin conformation assays including Hi-C. In the absence of chromatin conformation data, previously observed sizes of topologically associated domains (TADs) can be helpful as a prior.

We evaluated potential interactions between the identified promoter and enhancer elements. For this, both the spatial distance between the promoter and enhancer elements (up to 100 kb distance), as well as the correlation of expression (p-value < 0.05, Kendall’s tau > 0.3), were considered. We evaluated potential interactions between the identified promoter and enhancer elements by considering both spatial proximity (up to 100 kb) and correlated expression (p-value < 0.05, Kendall’s tau > 0.3). This yielded 11,298 potential enhancer–promoter interactions in the comprehensive set and 2,613 in the robust set. Most promoters were associated with only a few enhancers, whereas enhancers tended to interact with multiple promoters (Figure 6).

**Figure 6:**
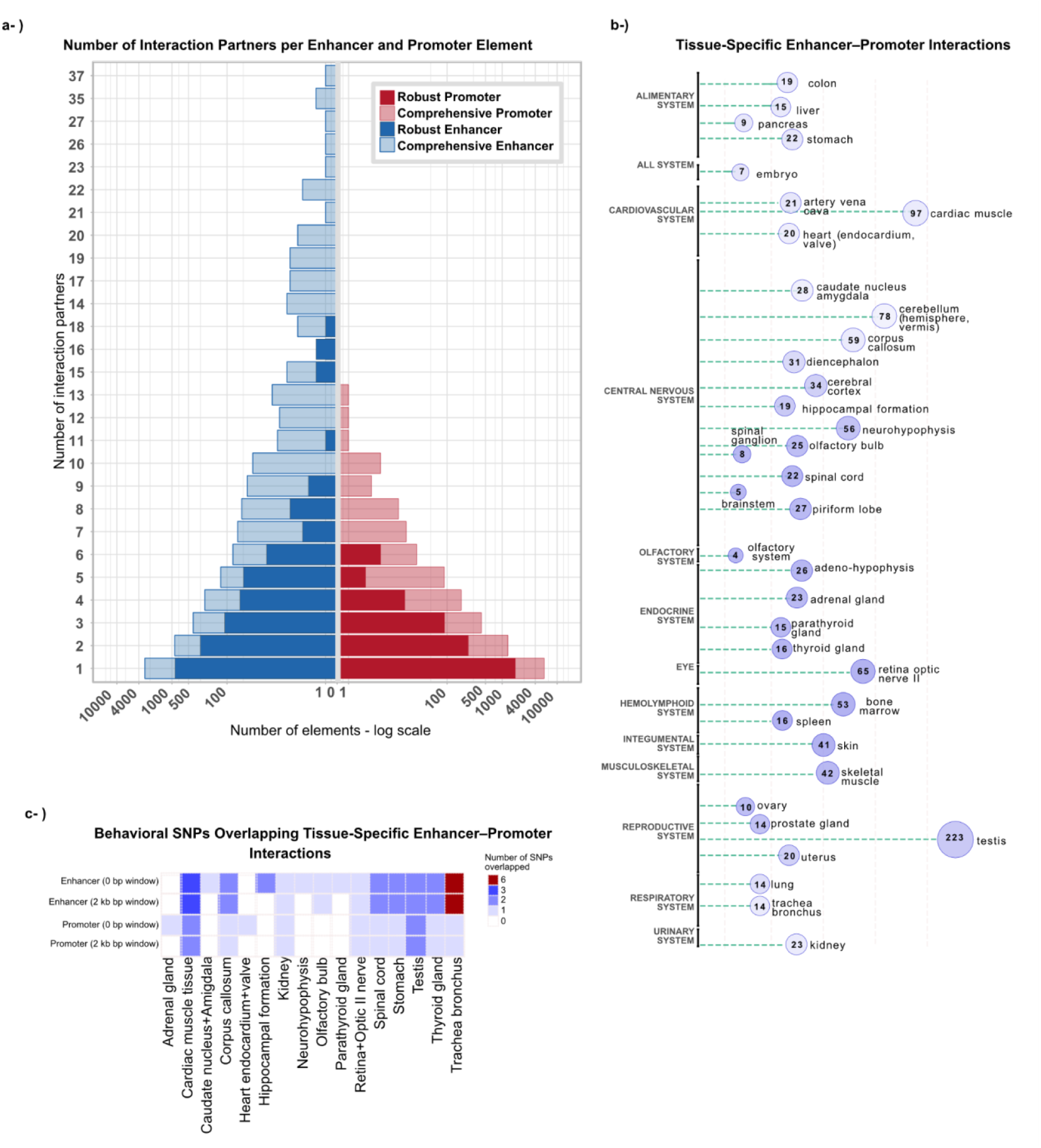
Enhancer-promoter interaction patterns: a) The figure illustrates the quantity of interaction partners. The bars on the left side of the figure indicate the number of interaction partners for the number of enhancers (the comprehensive set is represented by light yellow and the robust set by dark yellow), while promoters are depicted on the right (the comprehensive set is represented by light blue and the robust set by dark blue). b) The interactions between promoters and enhancers that occur within the same tissue are depicted. Numbers inside the circles indicate the number of interactions per tissue. c) The plot shows the number of behavioral SNPs that coincide in the interactions between promoters and enhancers that occur within the same tissue.

As promoters and enhancers were initially catalogued across bulk samples from multiple tissues to maximize element detection, not all inferred interactions were expected to be tissue-concordant. We therefore quantified the subset in which both linked elements were assigned to the same tissue. This subset comprised 1,221 enhancer–promoter interactions. Testis showed the highest number of same-tissue interactions (223), whereas the olfactory system showed the lowest. Within the central nervous system, the cerebellum had the highest number of same-tissue interactions (78).

### Enhancer and promoter activities are regulated by transcription factors

Genome-wide enhancer maps define tissue-specific regulatory landscapes but provide limited insight into the transcription factor architecture organizing regulatory activity. In particular, how enhancer and promoter regulatory programs are structured and coordinated across tissues remains poorly resolved at the organism scale. Leveraging the breadth of the DoGA dataset, spanning more than 100 tissues, we employed *de novo* motif discovery and network-based analyses to infer regulatory architectures underlying tissue-enriched enhancers and promoters.

We identified 1,247 *de novo* transcription factor binding site (TFBS) motifs within 700 bp (−500 to +200) of TSSs in tissue-enriched promoters and 970 *de novo* TFBS motifs within 400 bp (−200 to +200) centered on tissue-enriched enhancer midpoints. A substantial fraction of these motifs showed strong similarity to known transcription factor motifs in JASPAR 2024, with 757 promoter-associated and 565 enhancer-associated motifs exhibiting high concordance (Pearson correlation >0.9 and length-normalized correlation >0.5) (Figure 7, Tables S2-S3)

**Figure 7.**
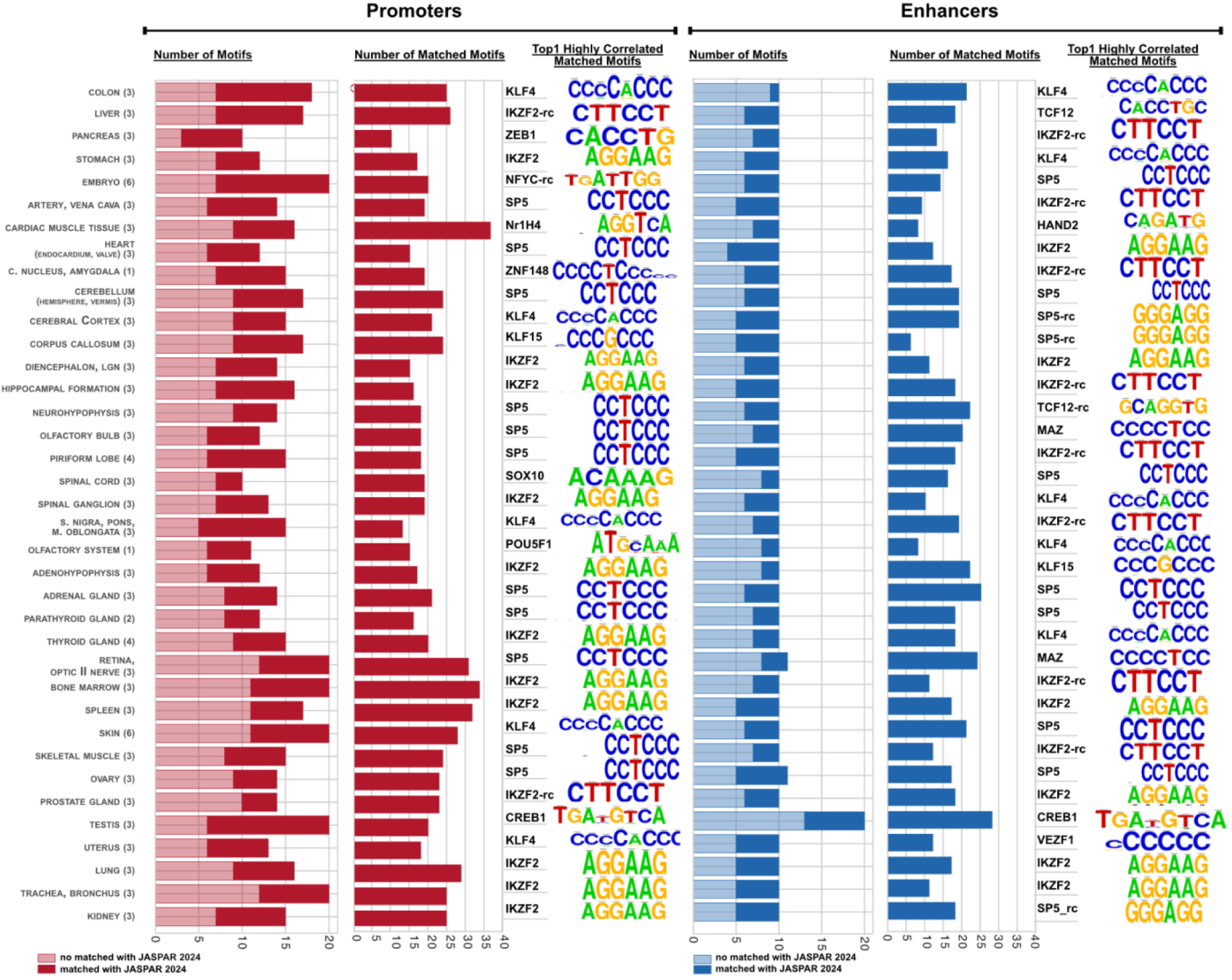
Distribution of *de novo* transcription factor binding site (TFBS) motifs and matched TFBS motifs. The figure displays the number of *de novo* motifs identified and their overlap within JASPAR2024. The top matched motifs for each tissue (cor > 0.9, n-cor > 0.5) are illustrated. The regions utilized for the analysis extended from original coordinates to 500 bp upstream and 200 bp downstream of TSS for promoters and 200 bp in both directions for enhancers.

To characterize regulatory organization, we constructed motif-based networks for enhancers and promoters (Table S4-S5). Both displayed dense intra-connectivity as well as extensive inter-connectivity, with more than 14 nodes with more than one connection (Figure 8, Figure S8, S9). Enhancer and promoter networks showed both shared and distinct transcription factor components. Several transcription factors were specific to enhancers or promoters, whereas others, including SP family members, MAZ, PATZ1, and multiple ZNF factors, were common to both regulatory element classes, indicating shared regulatory infrastructure.

**Figure 8:**
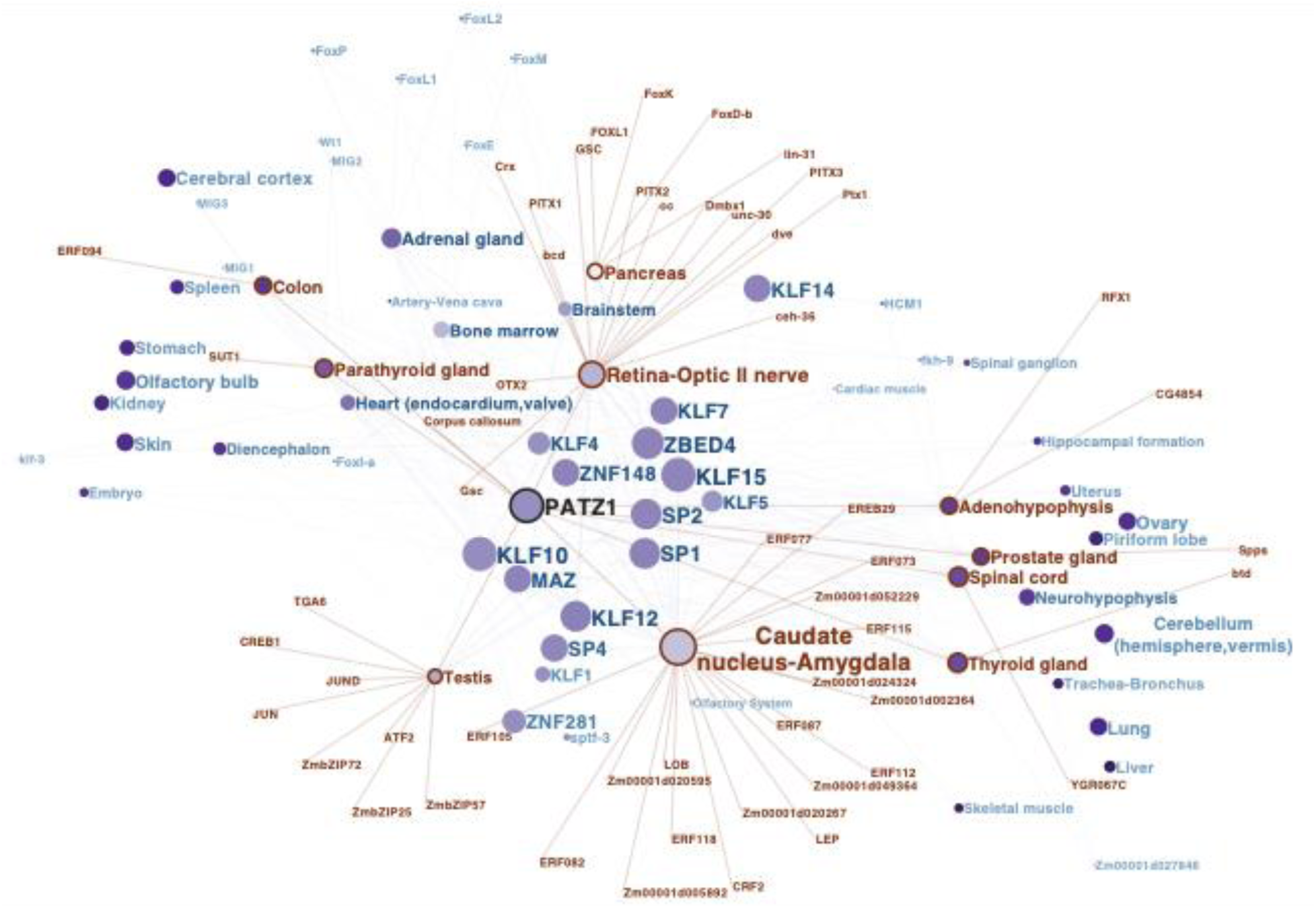
Combined enhancer-promoter motif network. The TFBS motif interaction network is constructed based on the intersection of enhancer and promoter motif networks (see individual enhancer and promoter networks in Figure S7, S8). Tissues are represented by red encircled nodes and red text color; region-specific TFBSs are depicted by red text; lighter and smaller text indicate less common connections. The nodes are positioned based on betweenness. The node size is proportional to their connectivity.

The KLF zinc finger transcription factor family members are located as an interconnected hub in the center of the promoter-enhancer intersection network. This positioning indicates their prevalence in nearly every tissue, thereby supporting their roles in diverse biological processes, including development, cell cycle regulation, and metabolism.

### Enhancer activity during early development in days 20-25 and 30

CAGE data from six embryos were selected to evaluate how embryonic enhancer activity develops during early development. In the course between days 20–25 and day 30, the activity of 54 enhancers was reduced, and the activity of 33 enhancers was increased on day 30 (Figure 9). To assess the functional roles of these enhancers, we took advantage of the potential enhancer-promoter interactions we established above. For the 54 down-regulated enhancers, we found regulatory interactions from 12 enhancers to 21 promoters, out of which 15 promoters were assigned to a known gene. For the 33 upregulated enhancers, we found that 8 enhancers possibly regulate 15 promoters, with 12 promoters corresponding to known genes. In total, there were 36 regulated promoters including 9 novel promoters, 7 primary promoters and 20 alternative promoters (Table S6-S8). Gene ontology enrichment analysis revealed that the regulated genes are involved in processes including neurodevelopment, early developmental regulation and chromatin-level control, genes shift to mature neuronal function, coupled with water-ion homeostasis and immune modulation.

**Figure 9:**
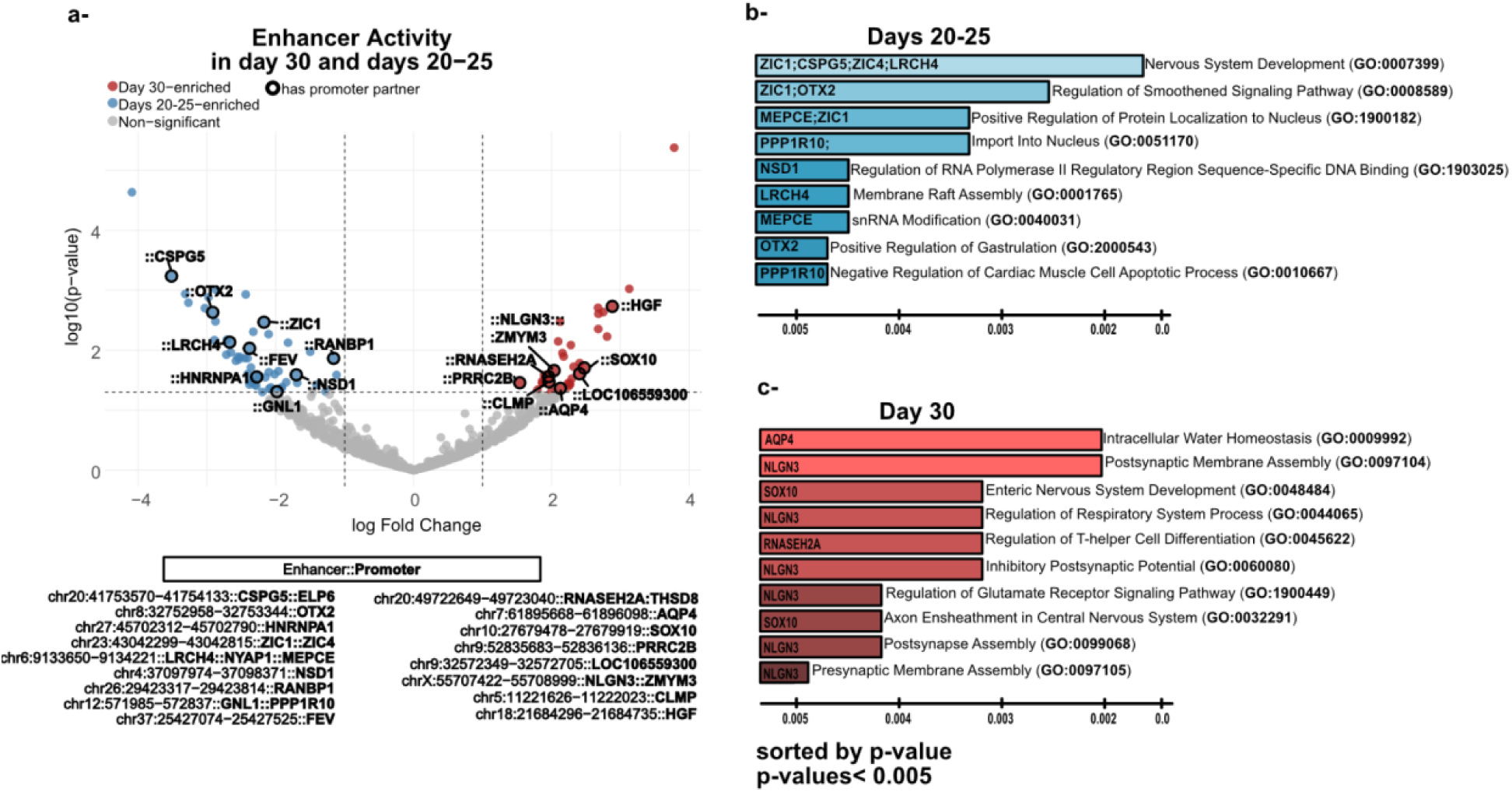
Use Case 1 - Embryonic enhancer activity occurs during days 20–25 and day 30. a) The active enhancers on day 30 are shown on the left with blue dots, while those on days 20–25 are on the right with red dots. The enhancers with a promoter partner are highlighted with a black circle. The gene name associated with that promoter is indicated at the end of the enhancer name in bold text in the table below and the only gene names are displayed on the plot. b) Gene ontology enrichment analysis for these genes that interact with the enhancers enriched in embryonic days 20–25 and c) day 30 stages.

### Enhancers and promoters are hotspots for genomic variants

Enhancers have been observed to harbor genomic variants with a frequency greater than expected by random chance^10^.

The promoter-enhancer interaction and the promoter-enhancer interaction partners enriched in the same tissue datasets were assessed to determine whether they coincide with any known SNPs. We found that approximately 3% of each dataset contains behavioral SNPs^24^ within the 2 kb window location of the elements and at least 75% of them within the exact coordinates of the element. “Howl” and “Toy directed Motor Patterns” associations are the most common SNPs among the promoters, while “Affectionate”, “Howls”, “Stares blankly”, “Dog Sociability” are most common for enhancers. Furthermore, nearly 4% of the promoter-enhancer interaction partners that were enriched in the same tissue coincided with behavioral SNPs (Table S8).

### Enhancers exhibit sequence similarity between dogs and humans

To understand sequence-level similarity of regulatory elements between dog and human, we compared 9,787 robust dog enhancers from our DoGA database with 65,423 human enhancers from the FANTOM5 database^25^ using BLAST^26^ dc-megablast to identify enhancer pairs sharing nucleotide similarity across species. We identified 1,312 dog enhancers that aligned with human enhancers. Of these, 1,199 enhancer pairs met our significance criterion (e-value < 1e-3).

We next examined 2,613 robust enhancer–promoter interactions derived from the distance (downstream 500 bp and upstream 1000 bp) and expression correlation of our CAGE promoters and enhancers. Intersecting these with the BLAST-aligned enhancer matches identified 403 dog enhancers that both (i) showed sequence similarity to a human enhancer and (ii) were involved in a dog enhancer–promoter interaction.

Based on these promoter–gene pairs, we ran an annotation-based mapping using Ensembl Compara with homology type “one2one.” From this set, we identified 93 potential human orthologs. GO enrichment analysis revealed significant functional signatures (adjusted p-value < 0.1) for Biological Process and Molecular Function. The annotated genes were enriched for developmental and tissue-specific biological process terms. Cellular component enrichment was not observed in this gene set.

From the original set of human enhancers, we collected the 1,199 enhancers that had a match from BLAST analysis. To map potential target genes for these human enhancers, we used the dataset from Engreitz Lab’s Activity-by-Contact (ABC) model-derived^27^. Human enhancers were intersected with ABC-predicted enhancer–gene pairs using BEDtools intersect, containing 8,092 human genes associated with these enhancers.

We then compared this set of ABC-derived human genes with the 93 human orthologs previously identified via Ensembl Compara from dog genes. Of the 8,092 ABC-derived human genes, 76 were also identified in one or more cell lines when compared to the 93 genes by Ensembl Compara.

## Discussion

We present the most comprehensive functional annotation of the canine genome to date. By applying CAGE-seq to 37 sample pools, representing 55 diverse tissues from 9 individual dogs, we generated 114 high-quality sequencing libraries to map the canine transcriptome and regulatory landscape. This effort identified 68,446 promoters, including 15,285 novel regions, and 46,661 active enhancers. By utilizing the CAGE technology, we move beyond histone-mark and open chromatin predictions to capture active transcription at the source^11^. While traditional ChIP-seq or ATAC-seq provides a map of regulatory potential, our detection of eRNA and TSS identifies the precise elements driving gene expression in real-time.

The technical robustness of these regulatory elements is validated by significant overlaps with H3K27ac and ATAC-seq marks. Specifically, 70% of our robust promoters and 46% of robust enhancers are supported by open chromatin peaks, while nearly half are marked by active acetylation. These findings, accessible through the DoGA Data Coordination Centre (DCC), provide a foundational resource that pushes the dog genome toward the gold standard annotation levels of human and mouse.

The study covers a wide range of tissues for promoter and enhancer discovery, but there is a particular interest in resolving the regulatory architecture of the canine brain, a critical step in using dogs as a spontaneous model for human mental disorders. Our dimension reduction analysis demonstrated that while most CNS samples group together, specific regions like the neurohypophysis and spinal ganglion maintain distinct regulatory signatures, reflecting their specialized neuroendocrine and sensory functions. Notably, the cerebellum emerged as a major regulatory hub, exhibiting 78 enriched enhancer-promoter interactions, the highest frequency observed among all brain regions in our dataset. This high level of connectivity likely reflects the cerebellum’s specialized requirement for fine-tuned motor coordination and its emerging role in higher-order cognitive and social behaviors^28–30^. By identifying these cerebellar-specific regulatory interactions, we provide a localized map for investigating the genetic basis of ataxia, broader neurodevelopmental phenotypes.

Beyond the brain, our analysis reveals distinct regulatory strategies across organ systems. The testis exhibited marked regulatory exceptionalism, harboring 3,214 enriched promoters and 1,177 robust enhancers. This pattern parallels classic observations that the testis displays unusually broad and permissive transcriptional activity compared with other tissues, including pervasive expression of lineage-restricted and young genes^31,32^. Rather than implying that the testis uniquely generates regulatory elements, our data are consistent with a model in which a permissive chromatin landscape and complex spermatogenic programs expose and activate a wide repertoire of promoters and enhancers, thereby increasing detectable regulatory diversity. In contrast, the eye exhibited a high-intensity, low-diversity profile. While it possessed a relatively small number of tissue-enriched enhancers, their average expression levels were among the highest in the study.

This suggests that while ocular development and maintenance rely on a more parsimonious set of regulatory elements compared to the testis, these enhancers operate with extreme transcriptional efficiency to maintain the physiological demands of the visual system^33^.

By integrating de novo motif discovery with network-based regulatory analysis, we show that canine regulatory programs are organized around shared transcriptional infrastructures. A prominent feature of this architecture is the enrichment of the KLF zinc finger family, which reflects the broader principle that transcription factor families often exhibit conserved DNA-binding specificities that enable coordinated regulatory control^34^. Within the inferred regulatory network, KLF15 displays high degree centrality, suggesting its potential role as a hub-associated regulator integrating signals across promoter- and enhancer-associated motifs. Rather than implying direct causal dominance, this observation is consistent with functional genomics studies showing that transcriptional homeostasis is often maintained by broadly connected regulatory factors, whereas tissue-specific regulation emerges from combinatorial enhancer motif assemblies.

The temporal resolution of the DoGA dataset allows for the dissection of the regulatory landscape occurring during canine embryogenesis. At days 20–25 of gestation, the enhancer landscape is dominated by a blueprint-level program targeting master regulators of neural tube morphogenesis and chromatin-level control, such as *OTX2*, *ZIC1/4*, and the epigenetic regulator *NSD1*^35,36^. By day 30, we observe a coordinated shift toward enhancers governing functional maturation. This includes the activation of enhancers linked to *AQP4* in brain water regulation and *NLGN3* in synaptic organization^37,38^. This transition from structural specification toward functional circuit–associated regulatory programs is consistent with a model in which canine embryogenesis recapitulates temporally organized neurodevelopmental regulatory dynamics relevant to human neurodevelopmental disorders, suggesting that the canine system may provide a valuable framework for studying vulnerability windows in neurodevelopment.

Our comparative analysis between 9,787 robust dog enhancers and the human FANTOM5 database reveals the characteristic rapid turnover of cis-regulatory elements^35^. Using BLAST-based alignment, we identified 1,199 dog enhancers with high sequence similarity to human enhancers. Crucially, when we intersected these with our enhancer-promoter interaction maps, we found 139 enhancers where both the sequence and the orthologous regulatory structure were conserved between species. This subset of 139 enhancers represents the core mammalian regulatory program. However, the majority of active dog enhancers lacked direct sequence homology to human elements, despite targeting orthologous genes. This suggests that regulatory function in dogs is often maintained through the conservation of transcription factor binding logic rather than primary nucleotide sequence, a phenomenon that has broad implications for translating canine genetic findings to human medicine.

While this study provides an unprecedented map of the canine regulatory landscape, several limitations remain. First, the relatively high RNA amount requirement for CAGE led to the pooling of certain tissues, which naturally results in a loss of ultra-fine tissue resolution in those specific libraries. Furthermore, our current comparative framework relies on sequence-based alignment, which often fails to detect elements that have undergone extensive primary sequence drift.

A promising future improvement involves the application of Interspecies Promoter-to-Promoter (IPP) synteny^39^. By using highly conserved promoters as syntenic anchors, IPP allows for the identification of regulatory elements based on their genomic neighborhood and relative position rather than sequence identity alone. As shown by Phan et al., many enhancers that appear species-specific due to sequence divergence actually retain deeply conserved regulatory functions across large evolutionary distances. Implementing such synteny-based models will likely reveal that a much larger fraction of the DoGA enhancerome is functionally conserved in humans, providing a more robust bridge for studying the non-coding variants underlying complex behavioral traits^24^.

In conclusion, the DoGA Consortium has established the most comprehensive atlas of active promoters and enhancers in the canine genome, significantly advancing our understanding of mammalian gene regulation. By resolving the spatial and temporal dynamics of the enhancerome from embryonic development to specialized adult tissues, we provide a high-resolution framework for using the dog as a spontaneous model for human disease. These data offer a transformative resource for the scientific community to explore the regulatory logic of morphological diversity, behavioral evolution, and the genetic architecture of multi-factorial traits.

## Methods

### CAGE library generation and sequencing

CAGE libraries were prepared following the previously described protocol by Takahashi et al.^40^. In total, 24 CAGE libraries were generated from 114 RNA samples. For library preparation, 1 μg of total RNA was taken from each RNA sample and the following barcodes were used to ensure that each sample within a library had a unique barcode: CTT, GAT, ACT, ATG, AGT, TGG, GTA, GCC, and ATC. In summary, total RNA was reverse transcribed to generate first-strand cDNA using random primers containing the EcoP15I sequence. Following reverse transcription, capped RNA-cDNA hybrids were selectively biotinylated at the 5’ cap and captured using the streptavidin-coated magnetic beads. Single-strand cDNA was then barcoded by adding a 5’ linker containing a 5’ barcode sequence and the EcoP15I sequence. After that, the second strand of the cDNA was produced. The resulting cDNA was then digested using the EcoP15I to cleave the cDNA at a sequence of 27 bp from the 5’ end. A 3′ linker with Illumina primer sequence was then added to the barcoded cDNA fragment. The resulting CAGE tags were then amplified and sequenced. The size of the resulting CAGE libraries ranged between 97 and 103 bp. Libraries were single-read sequenced at 50 cycles using the HiSeq 2500 Illumina System. The HiSeq SR Rapid Cluster Kit v2 (Illumina) and the HiSeq Rapid Duo cBot Sample Loading Kit (Illumina) were used to sequence CAGE libraries in the rapid mode.

### CAGE-seq data processing

All CAGE samples were processed by the nf-core/cageseq pipeline by modifying the development branch in nextflow DSL 2 (https://github.com/nf-core/cageseq/tree/dsl2)^41^. Following quality control using FastQC, trimming of adapters, EcoP15, and 5’G with cutadapt (the reads were kept even if the adapters or EcoP15, or 5’G were not detected) and rRNA filtering with SortMeRNA version 4.3.4 were conducted. Samples were aligned to CanFam4 using STAR^42^. Following an evaluation of paraclu and disclu clustering algorithms, the step involving CAGE tag counting and clustering in the nf-core pipeline was omitted, as the clustering performance was found to be most consistent with CAGEFightR^43^.

### CAGE tag cluster identification

The CTSS bigwig files from the nf-core workflow were imported into a RangedSummarizedExperiment (v1.20.0)^44^ object using CAGEfightR (v1.10.0)^43^. 33.16 million CTSSs were detected; 7.51 millions of them are supported by at least two samples.

### Promoter identification

The promoter regions were identified using the parameters of CAGEfightR’s ‘clusterUnidirectional()’ function, which applied pooled CAGE tags expressed at more than 10 tags per million (TPM) for the comprehensive set of promoters. The merge distances were set at 35 for the clustering. To provide a consistently and highly expressed robust subset, we kept only TSS with an expression of more than 10 tpm in at least two samples for the robust promoter set. This set of robust promoters consists of TSS from the comprehensive set, which are longer than 1 bp and meet the expression criteria. The promoter ranking was established based on TPM expression levels. The promoter with the highest TPM value was assigned p1, with subsequent promoters assigned in descending order of TPM, such as p2, p3, and so forth. The RefSeq gene model (assembly GCF_011100685.1)^22^ for the canine genome, referred to as canFam4, was utilized to annotate promoters, with a span of 500 bp downstream and 1000 bp upstream.

### Enhancer identification

Bidirectional clusters were identified with the default CAGEfightR parameters. Only candidates located in introns or intergenic regions were kept. Furthermore, enhancer candidates that overlapped with our robust promoter set were removed. For the comprehensive set, the candidates within the top 75% of all TPM normalized expression values were selected. Enhancers additionally had to have at least 4 different CAGE tag locations on each strand, and robust enhancers had to have at least 6 different CAGE tags on each strand. The tags that are present in the comprehensive set but absent in the robust set were evaluated. The tags with 98% bidirectionality are included in the robust enhancers. CAGEfightR sometimes produces regions that span wider than the CAGE tags. To compensate for that, we trimmed each enhancer region to the first and last CTSS within it.

### Promoters and enhancers are supported by epigenetic states

ATAC sequencing data provided by Barkbase were used^8^. We selected only those regulatory regions that contained at least one raw count from one of the tissues that were included in both studies. These tissues were bone marrow, liver, heart, stomach, spleen, thyroid, pancreas, skeletal muscle tissue, the pituitary gland and the cortex. Processing of the raw data was performed using the nf-core/atac-seq workflow. The cortex data were removed due to low quality and the number of peaks (Figure S16). The same overlapping was performed for the processed ChIP-seq peaks from the study by Son et al^9^. Epigenetic signal data were collected using coordinates obtained from CAGE-derived promoter and enhancer elements, which are both robust and comprehensive, to examine overlap. The mean signal value was determined for each sample and used as a threshold for each sample separately. If an element exceeded this threshold for the specific sample, it was classified as an overlap.

### Comparative genomics: promoter and enhancer conservation

PhyloP11 scores for the dog genome regions were obtained from the study published by Capriotti et al.^45^ based on alignments with 10 mammalian species (Human, Chimpanzee, Mouse, Rat, Cow, Panda, Marmoset, Cat, Horse and Opossum). We used regions of 400 bp around the midpoints of each element as input for deepTools (v3.5.1)^46^ to generate conservation plots. We generated 80,000 random regions of 400 bp (excluding enhancer, promoter regions and annotated Ensembl transcripts) using regioneR (v1.26.1)^47^. These regions acted as a reference for the background.

### Enhancer-promoter interaction

To obtain pairs of potentially interacting enhancers and promoters we applied a distance threshold of 100 kb.^27^ Then, we calculated Kendall correlation coefficients for the TPM normalized expression of all samples between the two elements that had to have a correlation of 𝛕 > 0.3 and a p-value of <0.05.

### Tissue enrichment scores

We calculated promoter and enhancer tissue-enrichment scores by taking the log fold change of the mean TPM value within one promoter or enhancer region for one tissue of interest, divided by the mean TPM value for each of the other tissues. We established three enrichment sets with 3, 5, and 10 thresholds for promoters and 3 and 5 sets for enhancers. To create those sets, we employed robust sets of promoters and enhancers. We selected a subset of these sets for downstream analysis. If the score was greater than 3, the promoter was defined as enriched in that tissue type for downstream analysis. For enhancers, the threshold was established at greater than three as well.

### Transcription factor binding site analysis

We looked for transcription factor binding sites around the tissue-enriched promoter and enhancer regions from the comprehensive data sets. First, we extended the promoters by 500 bp upstream and 200 bp downstream and the enhancers by 200 bp in both directions. We then identified the de novo motifs with RSAT’s peak-motifs function with default parameters^48^. The TRANSFAC format of the de novo motifs were utilized to evaluate similarity with known motifs from the JASPAR vertebrates, which were extracted from the JASPAR 2024^49^ core set, employing RSAT compare-matrices with a high stringency. This entailed a Pearson correlation coefficient exceeding 0.90 and a normalization with length exceeding 0.50, utilizing all identified promoter/enhancer regions as a background.

### Differential enhancer activity during development stages

Differential enhancer activity between embryonal stages 20-25 days versus 30 day was performed on robust enhancer raw count data. The total count across samples with fewer than 10 counts was filtered. Because enhancer RNAs typically exhibit low and variable read counts, we applied voomWithQualityWeights, which combines observational-level precision weights with sample-specific quality weights to improve variance modeling in sparse RNA-seq data^50,51^. Differentially active enhancers between developmental stages were identified using limma, and p-values were adjusted using the Benjamini-Hochberg method. Enhancers with nominal P < 0.05 and |log2 fold change| > 1 were classified as day 30-enriched or days20-25-enriched; all others were considered non-significant.

### Overlap with dog behavioral SNPs

We conducted an overlap analysis of comprehensive and robust enhancer and promoter sets with behavioral SNP clusters^24^. Morrill’s behavioral SNP clusters were collected, and their coordinates were converted from CanFam3.1 to CanFam4 with liftOver^52^. Redundancy resulting in some overlapping regions at the same location was removed. Interaction partners, enhancers and promoters, were split into two BED files as anchors. Duplicate anchors resulting from the many-to-many interaction relationship were removed from each file. Bedtools were employed to perform intersections on each set created subsequently, utilizing both a 2 kb window and without a window. This process involves creating BED files for promoters and enhancers and deduplicating them for co-enriched interaction partners as well.

### Investigation of the enhancer sequence similarity between dogs and humans

Robust dog enhancers were collected from CAGE-seq data. Human hg19 enhancers were downloaded from the FANTOM5 database^25^. To evaluate sequence-level similarity between dog and human enhancers, we performed pairwise sequence alignment using BLAST dc-megablast (2.16.0), which is optimized for detecting similar DNA sequences across species^26^. Each dog enhancer was queried against the human enhancer dataset using-max_hsps 1 to collect only the single best local alignment per enhancer pair. To ensure that only confidently aligned regions were retained, we filtered BLAST hits based on e-value < 1e-3. Robust enhancer–promoter interaction pairs from dog CAGE data were intersected with BLAST-aligned enhancer coordinates using BEDtools intersect with default parameters. For promoters involved in enhancer–promoter interaction pairs, we compared their genomic coordinates to an annotated promoter–gene set. Promoters with an exact match were assigned the corresponding annotated gene. Potential human orthologs for assigned genes were then retrieved through Ensembl Compara (Ensembl release 115)^53^. Genes with homology type of “one2one” were included for the next step. Gene Ontology enrichment analysis was performed by submitting a gene list to EnrichR^54,55^. Terms with adjusted p-values < 0.1 were considered significant. Top ten enriched terms from the Biological Process, Molecular Function, and Cellular Component ontologies were extracted and visualized using ggplot2. Human enhancers corresponding to the dog enhancer–gene pairs were further analyzed using a dataset derived by the Activity-by-Contact (ABC) model^27^ (Engreitz Lab; https://www.engreitzlab.org/resources) to predict potential enhancer–gene relationships. BED files for human enhancers were intersected with ABC-predicted enhancer–gene pairs using BEDtools intersect to collect 8,092 unique genes associated with these enhancers. These ABC-derived human genes were compared with the set of human orthologs identified by Ensembl Compara to assess overlap between sequence- and homology-based approaches.

## Author information

These authors contributed equally: Isil Takan, Matthias Hortenhuber

**Department of Medicine, Huddinge, Karolinska Institutet, Huddinge, Sweden** Işıl Takan, Matthias Hörtenhuber, Rijja Bokhari, Rasha Fahad Aljelaify, Faezeh Mottaghitalab, Amitha Raman, Fiona Ross, Carsten O. Daub & Juha Kere

**Department of Veterinary Biosciences, University of Helsinki, 00014, Helsinki, Finland** Marjo K. Hytönen, César L. Araujo, Ileana Quintero, Pernilla Syrjä, Noora Salokorpi, Antti Iivanainen, Ileana B. Quintero, Shintaro Katayama & Hannes Lohi

**Department of Medical and Clinical Genetics, University of Helsinki, 00014, Helsinki, Finland** Marjo K. Hytönen, César L. Araujo, Ileana Quintero, Noora Salokorpi, César L. Araujo, Hannes Lohi

**Folkhälsan Research Center, 00290, Helsinki, Finland** Marjo K. Hytönen, César L. Araujo, Ileana Quintero, Noora Salokorpi, César L. Araujo, Ileana Quintero, Sini Ezer, Juha Kere & Hannes Lohi

**Department of Equine and Small Animal Medicine, University of Helsinki, Helsinki, Finland** Tarja Jokinen

**Science for Life Laboratory, Karolinska Institutet, Stockholm, Sweden** Carsten Daub, Rasha Fahad Aljelaify, Matthias Hörtenhuber

**Department of Population Health and Reproduction, School of Veterinary Medicine, University of California, Davis, US** Danika Bannasch

**Roslin Institute and Royal (Dick) School of Veterinary Studies, University of Edinburgh, Edinburgh, Scotland** Jeffrey J. Schoenebeck

**Stem Cells and Metabolism Research Program, University of Helsinki, Helsinki, Finland** Sini Ezer, Shintaro Katayama & Juha Kere

## DoGA Consortium

Hannes Lohi, Juha Kere, Carsten O. Daub, Marjo K. Hytönen, César L. Araujo, Ileana B. Quintero, Kaisa Kyöstilä, Maria Kaukonen, Meharji Arumilli, Milla Salonen, Riika Sarviaho, Julia Niskanen, Sruthi Hundi, Jenni Puurunen, Sini Sulkama, Sini Karjalainen, Antti Sukura, Pernilla Syrjä, Niina Airas, Henna Pekkarinen, Ilona Kareinen, Anna Knuuttila, Hanna-Maaria Javela, Laura Tuomisto, Heli Nordgren, Karoliina Hagner, Tarja Jokinen, Antti Iivanainen, Kaarel Krjutskov, Sini Ezer, Shintaro Katayama, Masahito Yoshihara, Auli Saarinen, Abdul Kadir Mukarram, Matthias Hörtenhuber, Rasha Fahad Aljelaify, Fiona Ross, Faezeh Mottaghitalab, Işıl Takan, Noora Salokorpi, Amitha Raman, Irene Stevens, Oleg Gusev, Danika Bannasch, Jeffrey J. Schoenebeck, Heini Niinimäki & Marko Haapakoski, Minna E. Ikonen

## Contribution statements

HL, MKH, PS recruited the animals. IQ performed RNA extraction. AR and RA generated CAGE libraries and data. DCC was built by CA. Data analysis was performed by IT, MH, NS, MEI, FR, FM, SE, SK and AI. Data management was conducted by RB. Study was conceived, planned and supervised by HL, JK and CD.

## Acknowledgments

Sini Karjalainen is thanked for technical assistance. CSC is thanked for data hosting and computing resources. The computations were enabled by resources in project sens2020609 provided by the National Academic Infrastructure for Supercomputing in Sweden (NAISS) at UPPMAX, funded by the Swedish Research Council through grant agreement no. 2022-06725. We thank the Centre for Bioinformatics and Biostatistics (CBB) at Karolinska Institutet for providing computational resources. The study was funded in large part by the Jane and Aatos Erkko Foundation and partially by Science & Diagnostics Unit, Mars Petcare.

## Data availability

All data are available from the DoGA Data Coordination Centre under https://dcc.doggenomeannotation.org/doga/data_export_worker/.

All sequences of the CAGE samples and the most processed data of these sequences will be available in NCBI’s GEO Accession GSE313038.

Computational workflows are accessible in the DoGA Git repository under https://gitlab.com/doggenomeannotation/.

An online expression atlas to explore gene, promoter, and expression can be accessed at https://expression-atlas.doggenomeannotation.org/dogaatlas/.

An example of the Zenbu view is available at https://fantom.gsc.riken.jp/zenbu/gLyphs/#config=c8zMMypAD5wt_UekszItgB;loc=canFam4::chr20:57977094..58039708+

