## Supplemental Figures S1-S16 for "The DoGA Consortium Atlas of Canine Enhancers and Promoters Across Tissues and Development"

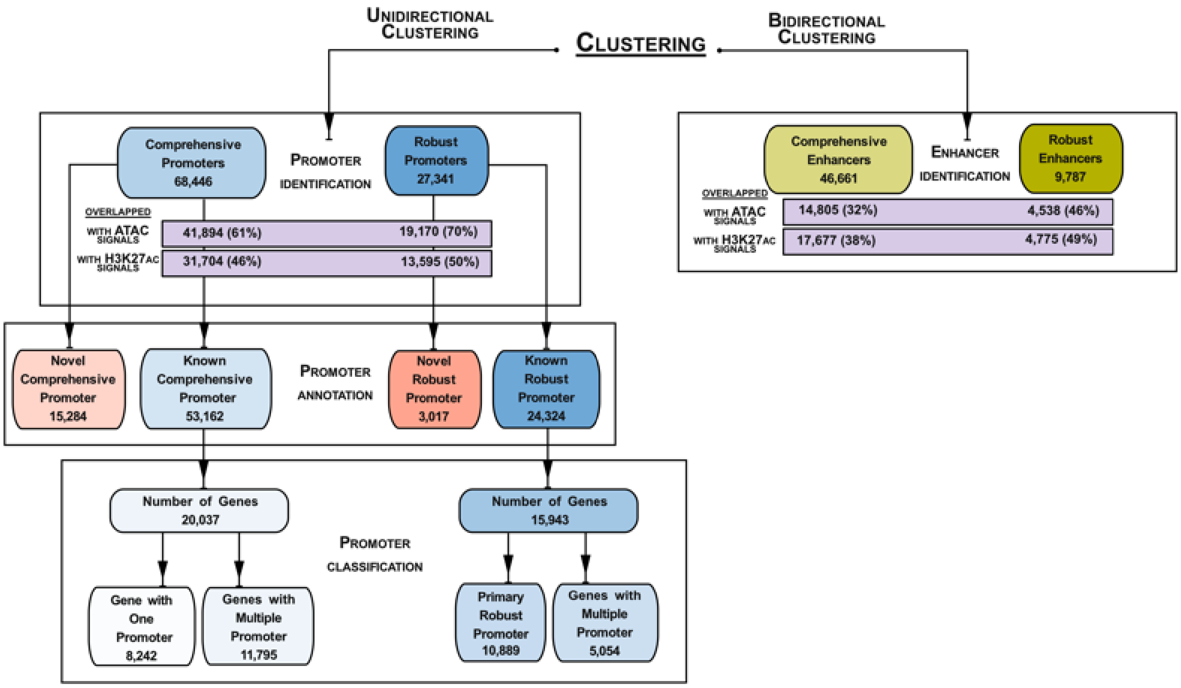


Figure S1: The number of regulatory elements (promoters and enhancers).

The 'promoter identification' box shows the number of promoters identified with external validation. We established comprehensive and robust collections of promoter elements based on expression level, the number of samples exhibiting that level, and the width of the element.
The number of elements overlapped with ATAC signal and H3K27ac signals was displayed. Percentages represent the total number of epigenetic signals relative to the overlapped signal for each marker. The 'promoter annotation' box shows the number of annotated and novel to CanFam4 for both sets. The 'promoter classification' box shows the promoter-gene relations. We classified promoters as genes with one promoter and genes with multiple promoters. The 'enhancer identification' box shows the number of enhancers identified with external validation.

| | Alaskan Malamute | Border Collie | Collie | East-European Shepherd | Finnish Lapphund | German Pinscher | Mixed Breed | Rottweiler | Swedish Elkhound |
|---|---|---|---|---|---|---|---|---|---|
| Colon | | | | | 1 | 1 | | 1 | |
| Liver | | | | | | 1 | | 1 | 1 |
| Pancreas | | 1 | | | 1 | | | 1 | |
| Stomach | | 1 | | | | 1 | | | 1 |
| Embryo | | | | | | | 6 | | |
| Artery, Vena Cava | 1 | | | | | 1 | | 1 | |
| Cardiac Muscle Tissue | | | | | | 1 | | 1 | 1 |
| Heart (Endocardium, Valve) | | | 1 | 1 | | 1 | | | |
| Caudate Nucleus,amygdala | | | | | | | | | 1 |
| Cerebellum (Hemisphere, Vermis) | | | | | 1 | | | 1 | 1 |
| Cerebral Cortex | | | | | 1 | | | 2 | |
| Corpus Callosum | | 1 | | | | 1 | | 1 | |
| Diencephalon, LGN | | 1 | | | 1 | | | 1 | |
| Hippocampal Formation | | | | | | 1 | | 2 | |
| Neurohypophysis | | | | | | 1 | | 1 | 1 |
| Olfactory Bulb | | 1 | | | 1 | | | | 1 |
| Piriform Lobe | | | | | 1 | | | 1 | 2 |
| Spinal Cord | | 1 | | | | 1 | | | 1 |
| Spinal Ganglion | | 1 | | | 1 | | | 1 | |
| Brainstem | | 1 | | | | | | 1 | 1 |
| Adenohypophysis | | | | | | 1 | | 1 | 1 |
| Adrenal Gland | | | | | 1 | | | 1 | 1 |
| Parathyroid Gland | | | 1 | | | 1 | | | |
| Thyroid Gland | | 1 | | | | 1 | | 1 | 1 |
| Retina, Optic II Nerve | | | | | | 1 | | 1 | 1 |
| Bone Marrow | | | | | 1 | | | 1 | 1 |
| Spleen | | | | | 1 | 1 | | 1 | |
| Skin | | | | | | 2 | | 4 | |
| Skeletal Muscle | | | | | | | | 2 | 1 |
| Olfactory System | | | | | | | | 1 | |
| Ovary | 1 | | | | | 1 | | | 1 |
| Prostate Gland | | 1 | | | | | | 2 | |
| Testis | | | 1 | | | | | 2 | |
| Uterus | | | | 1 | | 1 | | | 1 |
| Lung | | | | | | 1 | | 1 | 1 |
| Trachea, Bronchus | | | 1 | | 1 | | | 1 | |
| Kidney | | 1 | | | | 1 | | | 1 |

Figure S2: Sample collection with details.

CAGE tissues and pools, are listed alphabetically. Each column represents a different breed. Each tissue row indicates the number of breeds present in that tissue.

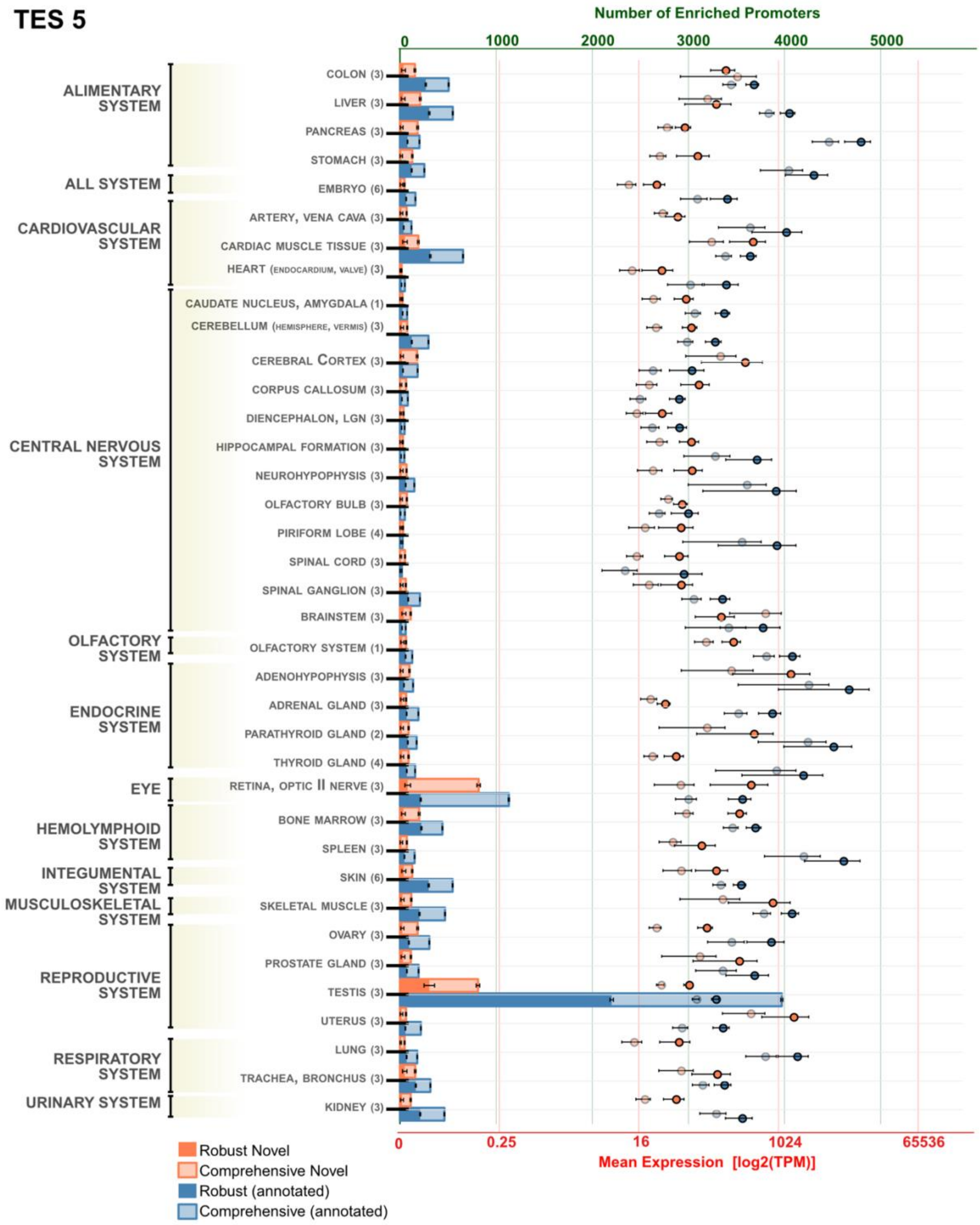


Figure S3: Tissue Enrichment Score 5 - Promoter.

The figure shows promoters with a tissue enrichment score of five for each tissue. The y-axis shows CAGE tissue pools and the related organ systems, listed alphabetically. The top x-axis indicates the number of promoters enriched in each tissue, while the bottom x-axis shows the mean expression (log2 TPM) of elements enriched in those tissues.

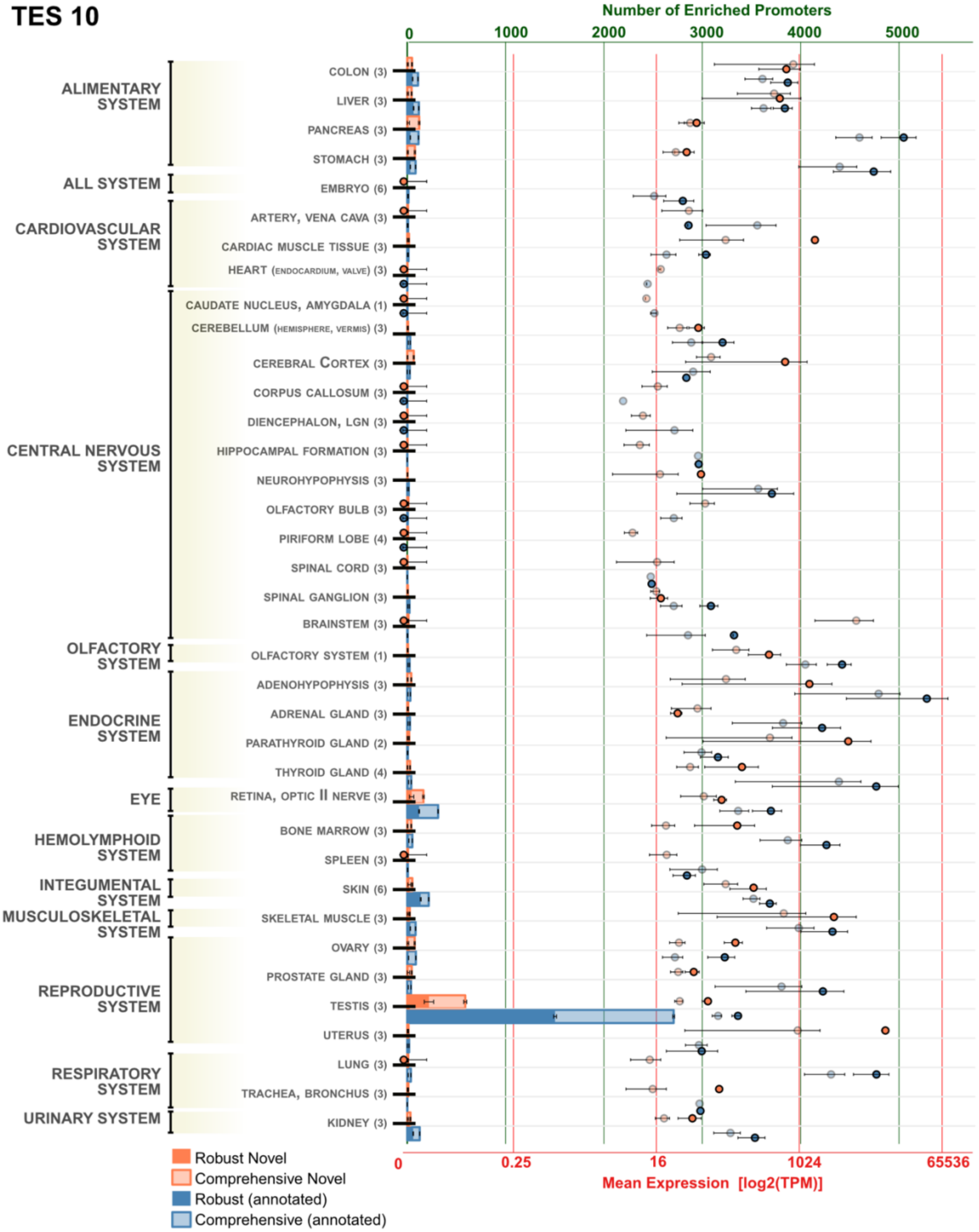


Figure S4: Tissue Enrichment Score 10 - Promoter.

The figure shows promoters with a tissue enrichment score of ten for each tissue. The y-axis shows CAGE tissue pools and the related organ systems, listed alphabetically. The top x-axis indicates the number of promoters enriched in each tissue, while the bottom x-axis shows the mean expression (TPM) with log2 scale of elements enriched in those tissues.

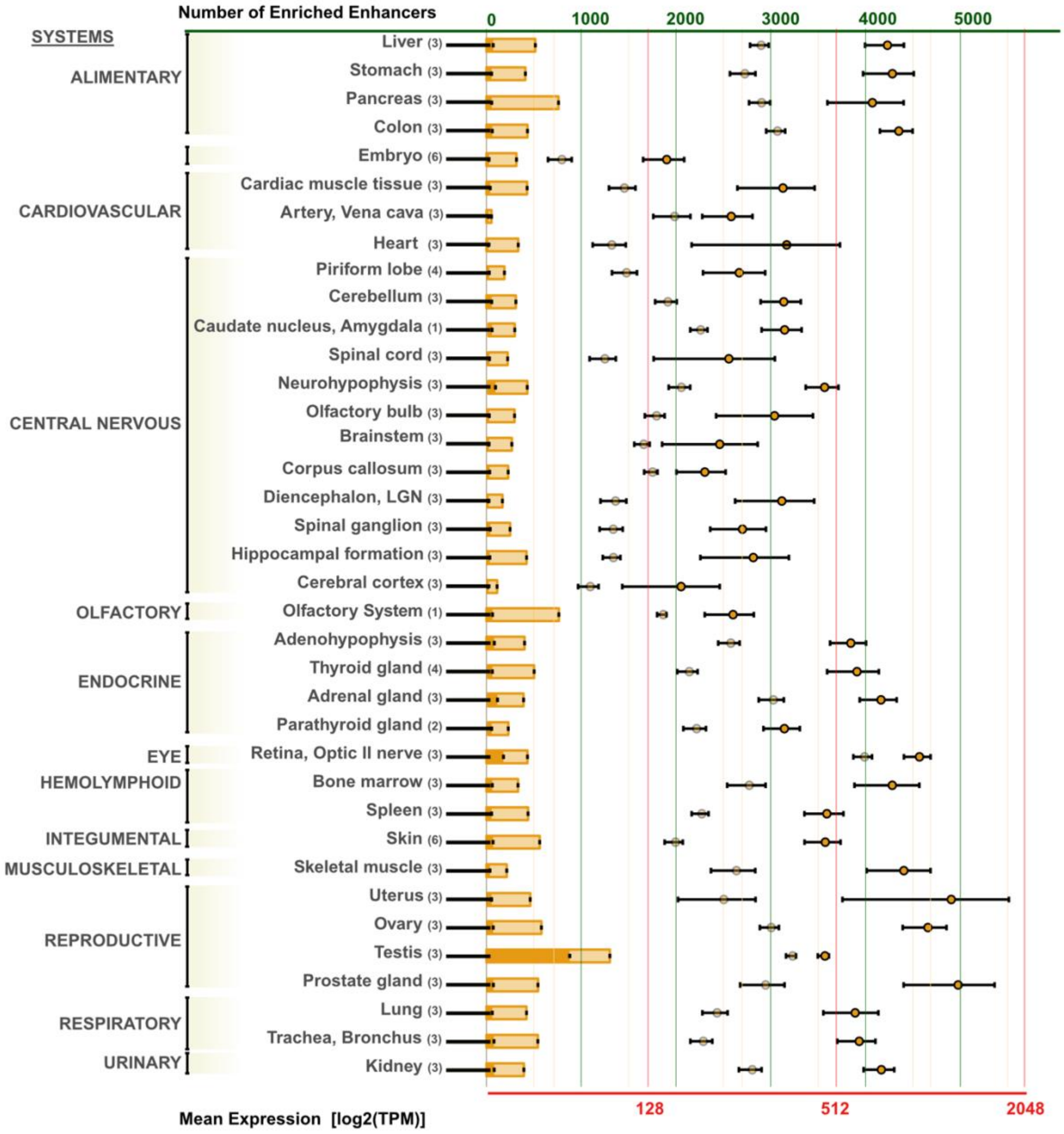


Figure S5: Tissue Enrichment Score 5 - Enhancer.

The figure shows enhancers with a tissue enrichment score of ten for each tissue. The y-axis shows CAGE tissue pools and the related organ systems, listed alphabetically. The top x-axis indicates the number of enhancers enriched in each tissue, while the bottom x-axis shows the mean expression (TPM) with log2 scale of elements enriched in those tissues.

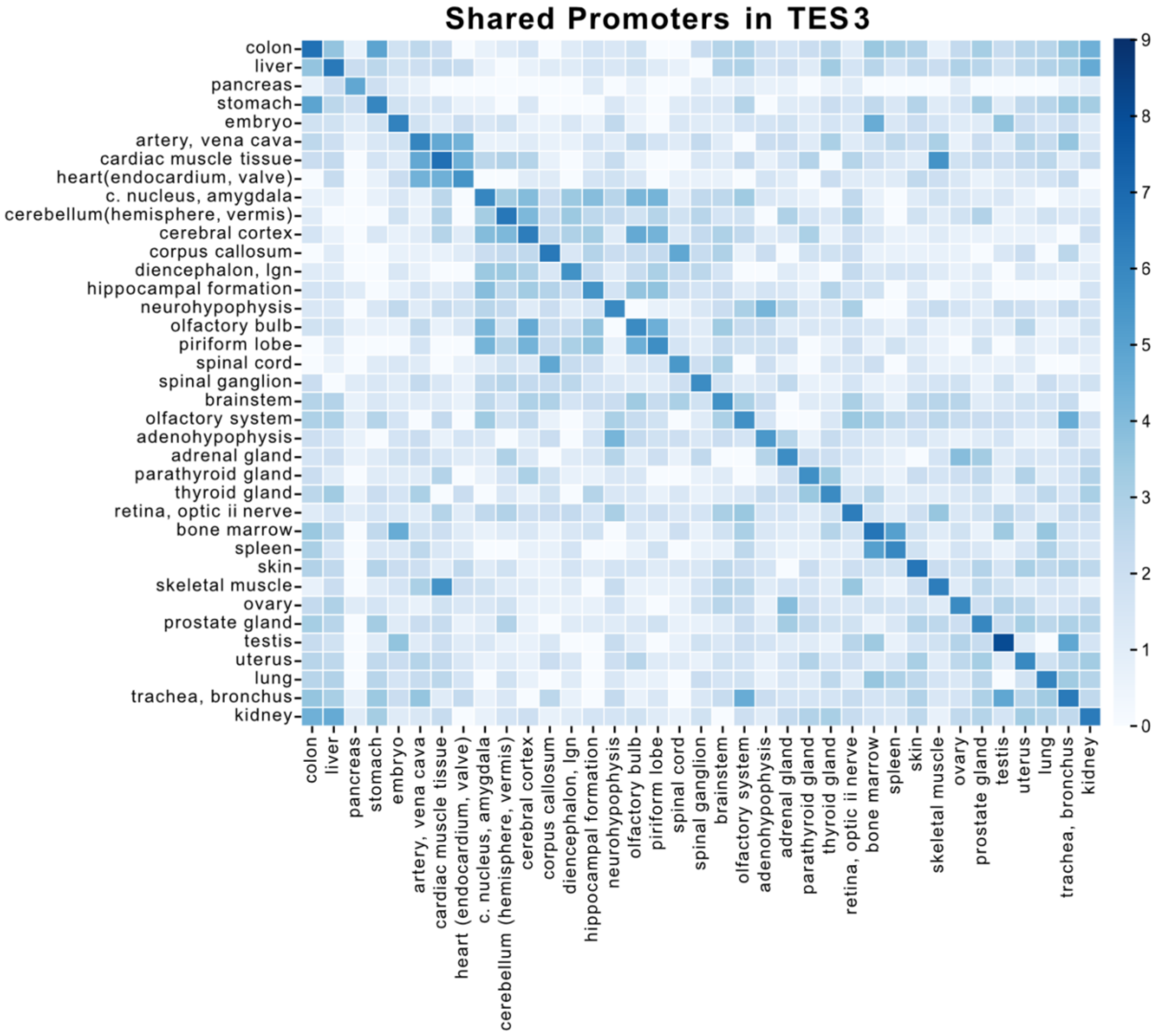


Figure S6: The common promoters in Tissue Enrichment Score 3.

The heatmap displays the common promoters with the log-transformed numbers in tissue enrichment. The matrix indicates the comparison of each promoter element enriched in tissues (TES 3 enriched promoter set was used), indicating common promoters between tissue pairs.

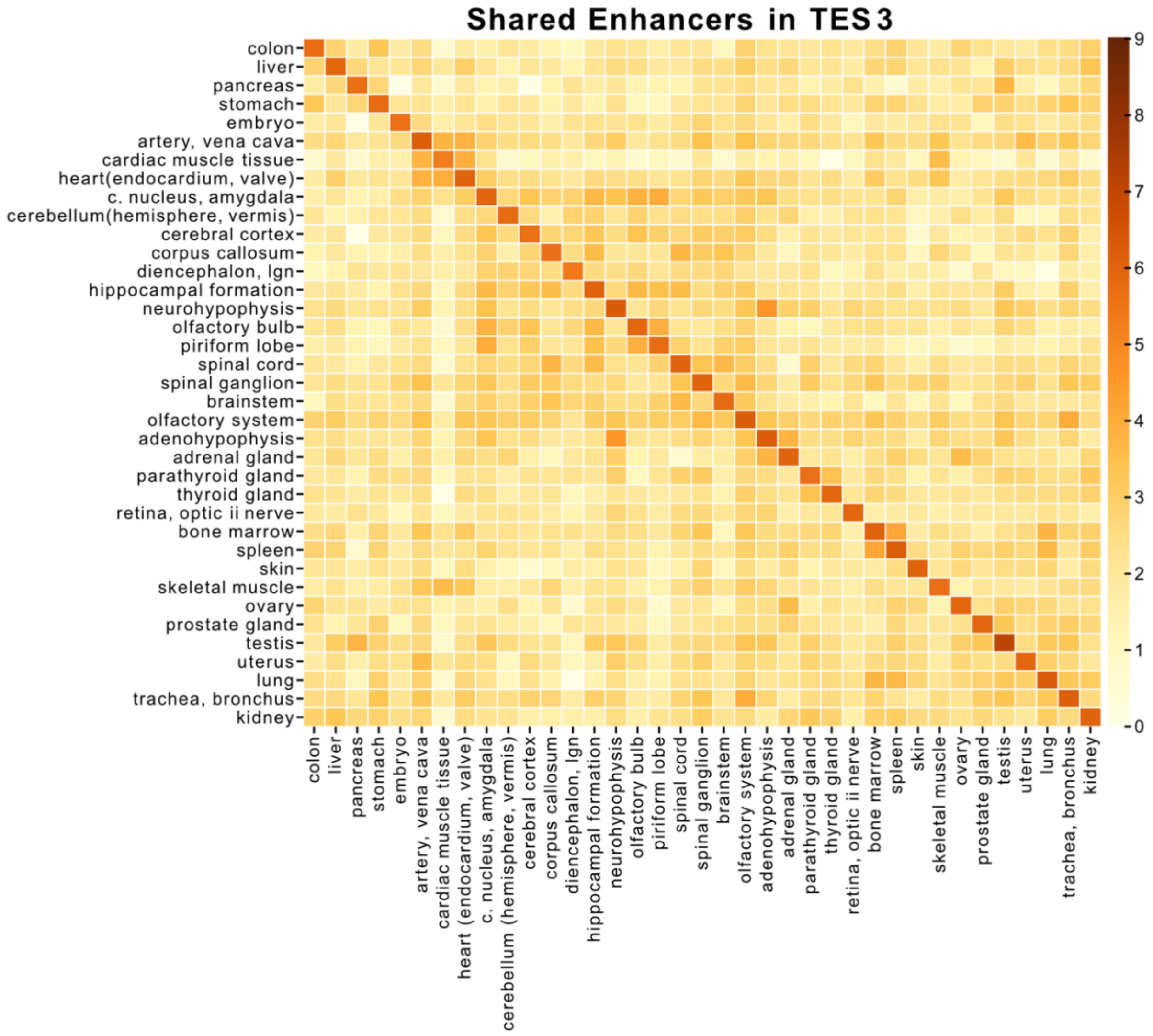


Figure S7: The common enhancers in Tissue Enrichment Score 3.

The heatmap displays the common enhancers with the log-transformed numbers in tissue enrichment. The matrix indicates the comparison of each enhancer element enriched in tissues (TES 3 enriched enhancer set was used), indicating common enhancers between tissue pairs.

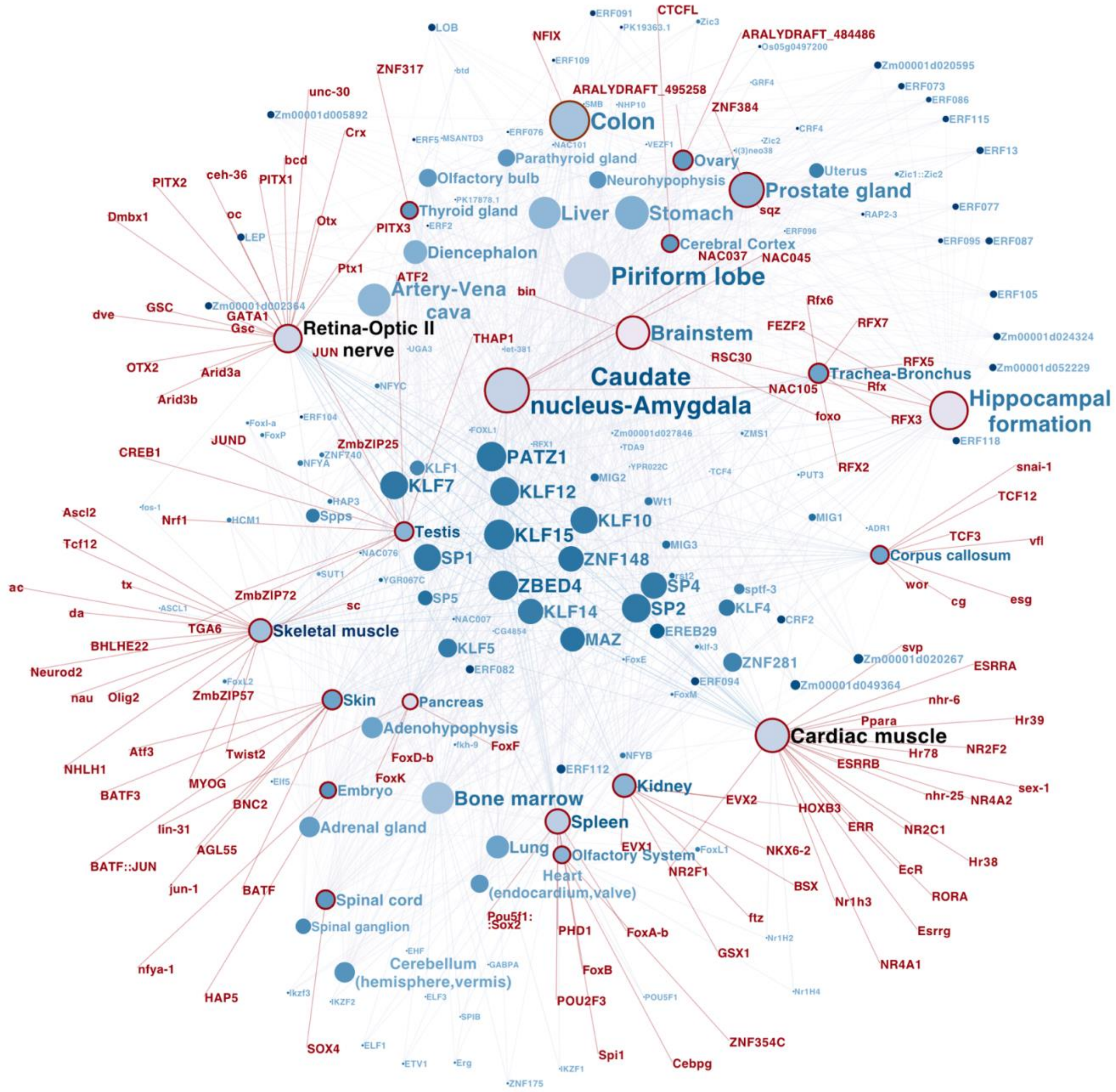


Figure S8: TFBS enrichment network (promoter)

The TFBS motif interaction network illustrates the distribution of the transcription factor motifs identified in promoter sequences across all tissues. The network is constructed based on the known motif (JASPAR 2024), which is enriched in the sequences of our promoter regions with high concordance (Pearson correlation >0.9 and length-normalized correlation >0.5). Tissues are indicated by red encircled nodes; region-specific TFBS are depicted by red text; lighter and smaller text indicate less common connections. The nodes are positioned based on Betweenness. The node size is proportional to their connectivity.

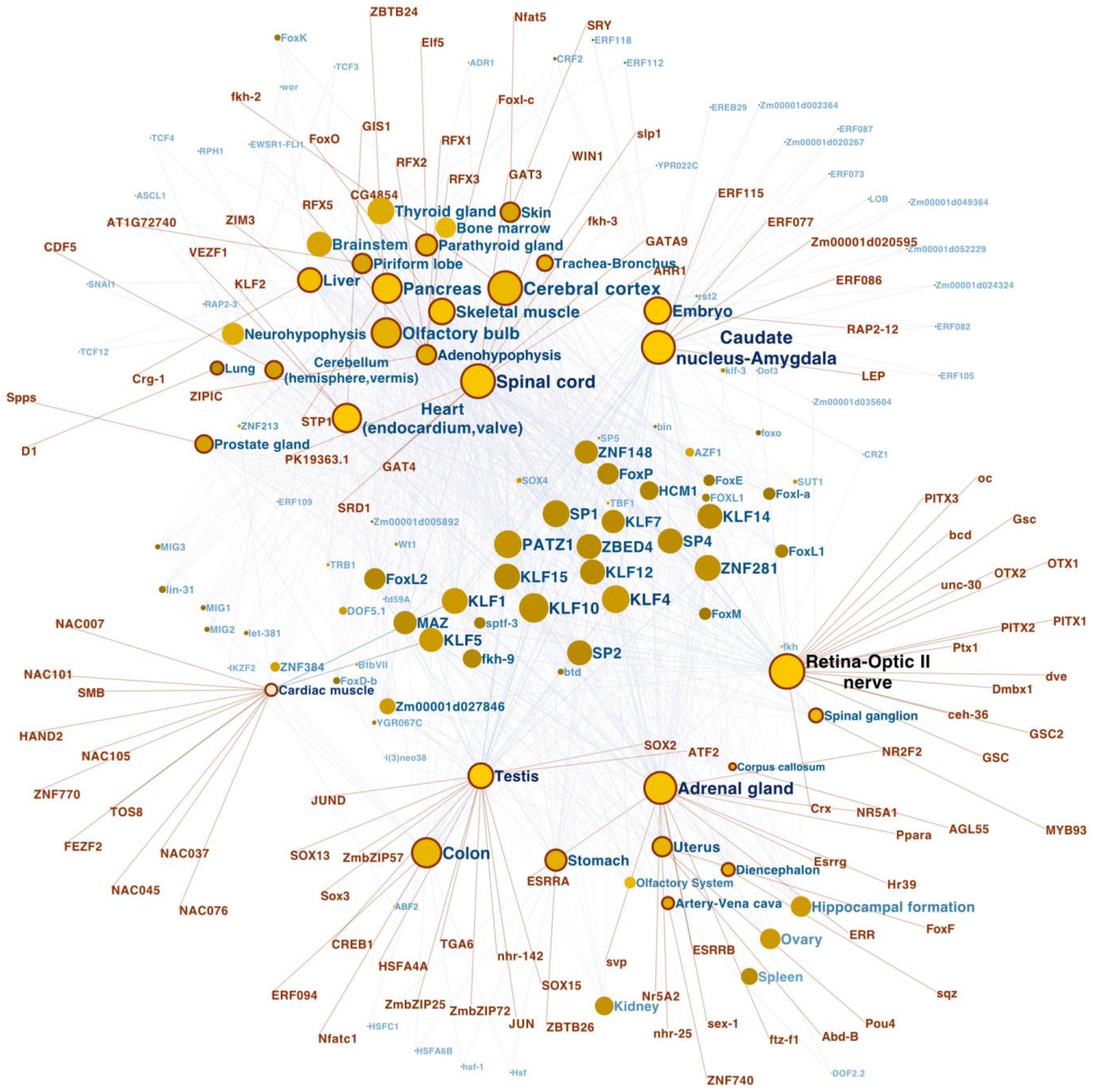


Figure S9: TFBS enrichment network (enhancer)

The TFBS motif interaction network illustrates the distribution of the transcription factor motifs identified in enhancer sequences across all tissues. The network is constructed based on the known motif (JASPAR 2024), which is enriched in the sequences of our enhancer regions with high concordance (Pearson correlation >0.9 and length-normalized correlation >0.5). Tissues are indicated by red encircled nodes; region-specific TFBS are depicted by red text; lighter and smaller text indicate less common connections. The nodes are positioned based on Betweenness. The node size is proportional to their connectivity.

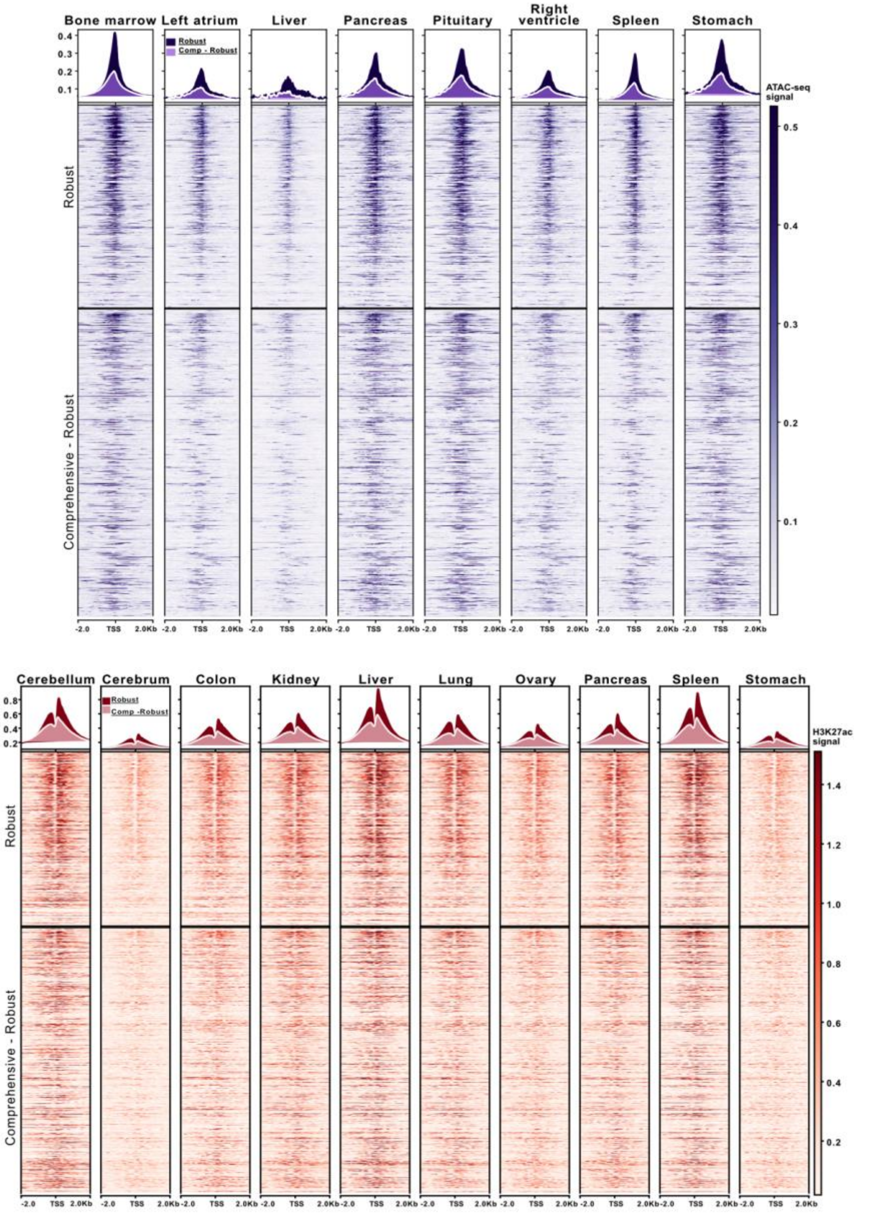


Figure S10: Evaluation of the DoGA promoter candidates by open chromatin marks and histone marks.

The heatmaps show external validation of the promoter candidates by open chromatin and H3K27ac marker. (A) Eight different dog tissue samples for ATAC-seq and (B) eleven for H3K27ac ChIP-seq data were analyzed. All heatmaps are sorted by expression of CAGE-defined promoters. The range of heatmaps spans +/- 2000 bp around the midpoints of each promoter. The line plots show mean values for each signal column. Strong signals for promoters were observed in ATAC-seq data, and a typical peak-valley-peak pattern appeared in H3K27ac ChIP-seq data.

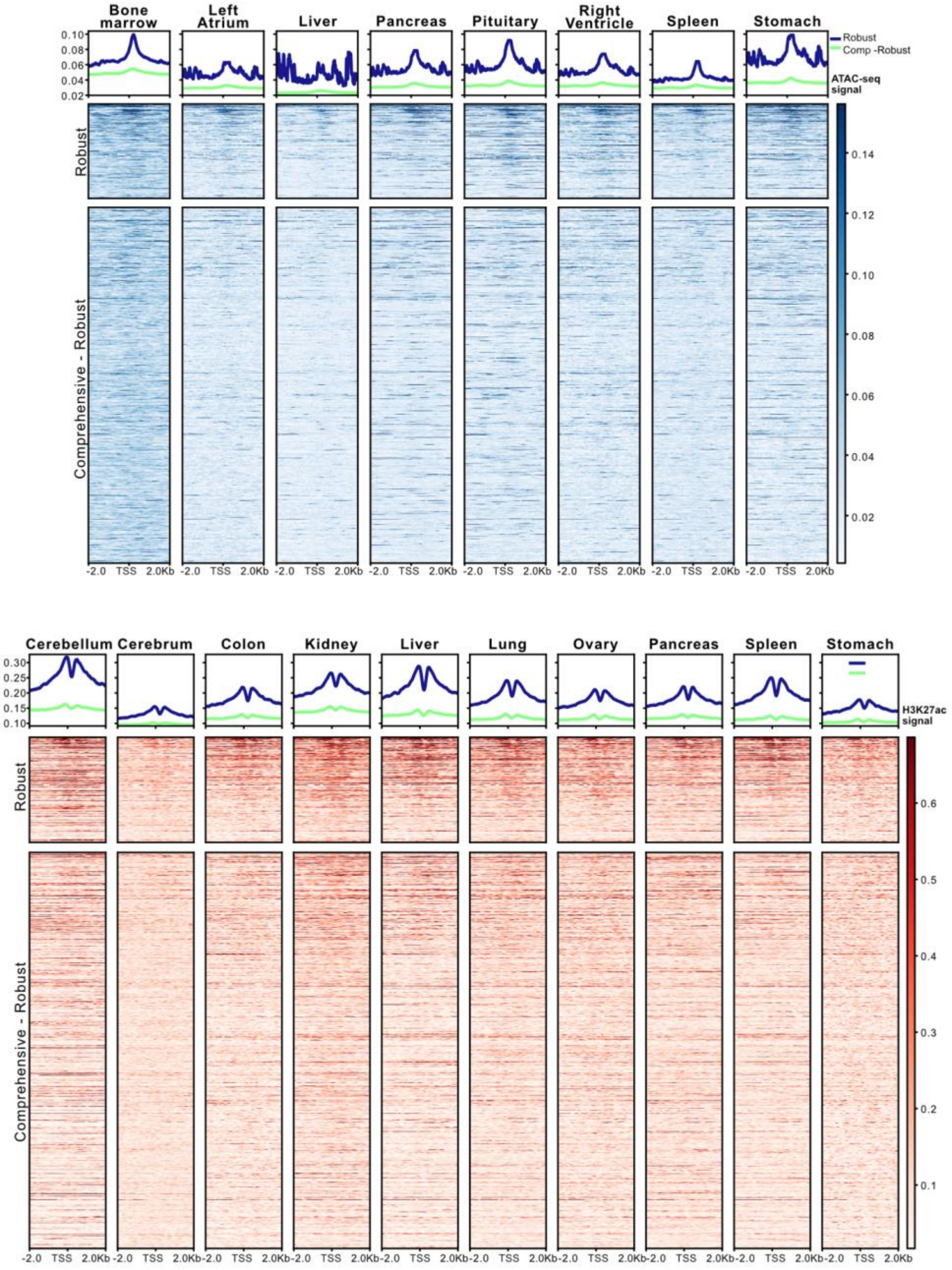


Figure S11: Evaluation of the DoGA enhancer candidates by open chromatin marks and histone marks.

The heatmaps show external validation of the enhancer candidates by open chromatin and H3K27ac marker. (A) Eight different dog tissue samples for ATAC-seq and (B) eleven for H3K27ac ChIP-seq data were analyzed. All heatmaps are sorted by expression of CAGE-defined enhancers. The range of heatmaps spans +/- 2000 bp around the midpoints of each enhancer. The line plots show mean values for each signal column. Notable peaks with relatively weak signals compared to promoters were observed in ATAC-seq data, and a typical peak-valley-peak pattern appeared in H3K27ac ChIP-seq data.

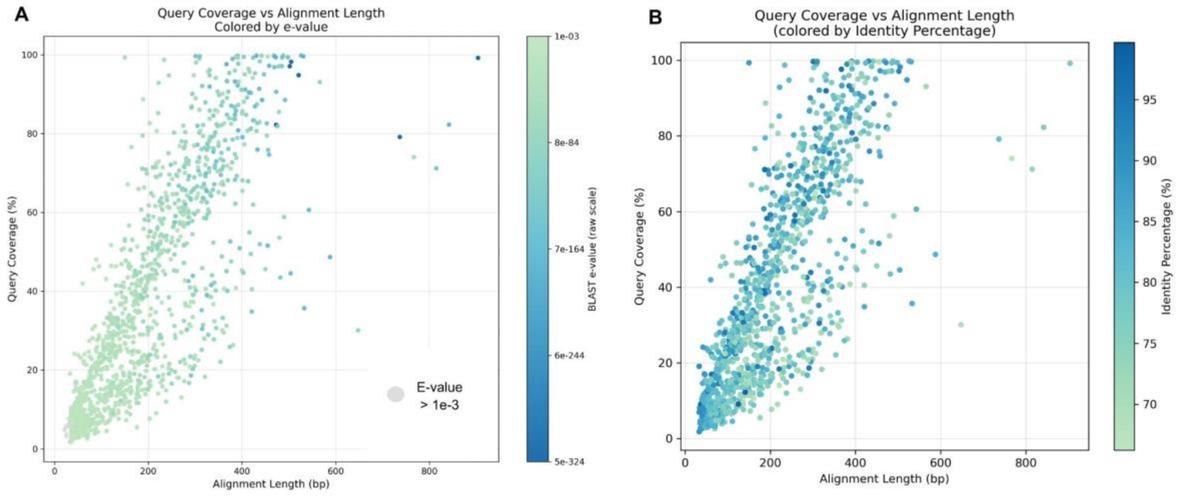


Figure S12: Comparison of BLAST alignment results between query sequences (DOGA dog enhancers) and subject sequences (FANTOM5 human enhancers).

(A) points are coloured according to BLAST e-value, with darker colours indicating more significant matches. (B) points are coloured by identity percentage, with higher identity values shown in darker colours. In both figures, each point represents a single BLAST match, with alignment length (bp) shown on the y-axis and percentage of query coverage on the x-axis.

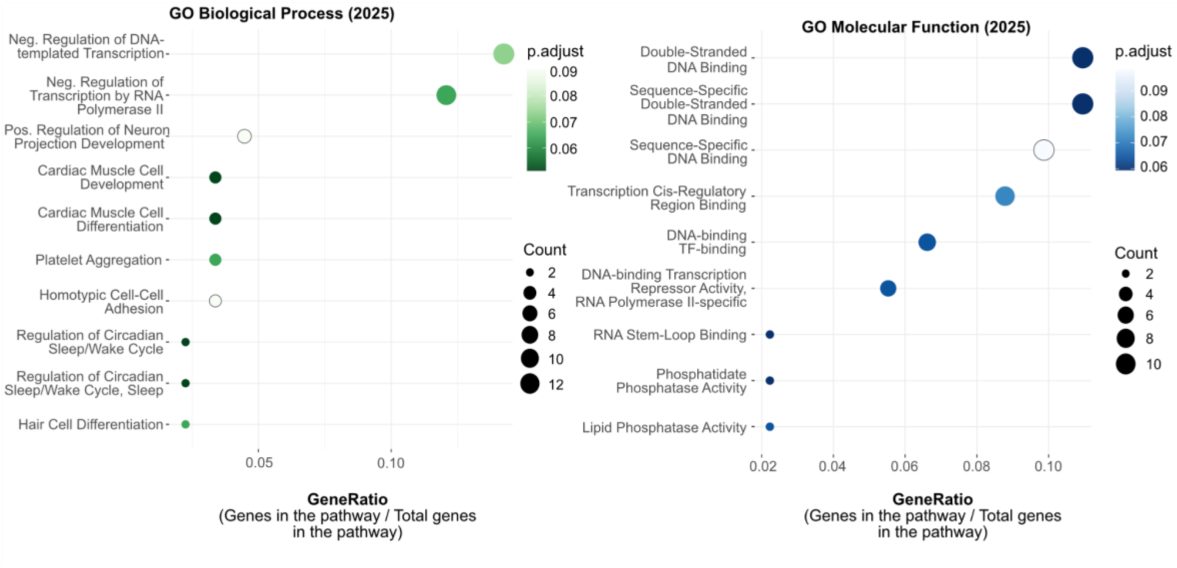


Figure S13: Gene ontology enrichment analysis of potential human orthologs using EnrichR.

Enriched Gene Ontology (GO) terms for the Biological Process and Molecular Function ontologies were illustrated, while Cellular Components didn't have any ontologies exceeding the set limit value. The x-axis represents the gene ratio (the proportion of input genes annotated to each GO term), while the y-axis lists the enriched GO terms. The size of each dot corresponds to the number of genes associated with each term, and the color reflects the adjusted P value (Benjamini–Hochberg FDR). Only terms that meet the enrichment cutoff (<0.1) are displayed.

Query dog robust enhancer set: 9,787 enhancers
Subject human permissive enhancer set: 65,423 enhancers
Enhancers were collected from all tissues available.

**Step 1**. Enhancer sequences from dog and human were compared based on nucleotide sequence similarity using BLAST (dc-megablast) with parameter --max-hsps=1, collecting only the best match for each human sequence based on expect value. Altogether, 1,312 matches were found.

**Step 2.** Enhancer matches were filtered to remove low-confidence alignments, excluding matches with e-values above 1e-3. 1,199 matches passed the filtering criteria.

**Step 3.** Dog enhancers with human sequence similarity were intersected with an existing robust enhancer–promoter dataset (2,613 pairs) using BEDTools intersect, considering only direct overlaps and no windowing, to collect associated promoters. We found 403 overlapping enhancers.

Robust annotated set contained promoters with gene annotation, within with a range of 500 bp downstream and 1000 bp upstream. This set consisted of 139 promoter with gene annotation.

**Step 4.** The genes corresponding to the promoters were extracted, and for multiple promoters of the same gene, only unique gene names were kept. The dog genes were mapped based on annotation to human genes using Ensembl Compara. Only genes with homology type "one2one" were included while "one2many" were excluded to to avoid genes with multiple possible mapped genes.

Robust annotated set: 93 potential orthologs

**Step 5.** Potential human orthologs were mapped to enhancers regulating the specific genes, based on ABC Model. The enhancers coordinates were compared to 1199 original subject enhancers from BLAST analysis with BEDtools intersect.

We found 76 same genes from ABC-derived genes and potential orthologs from

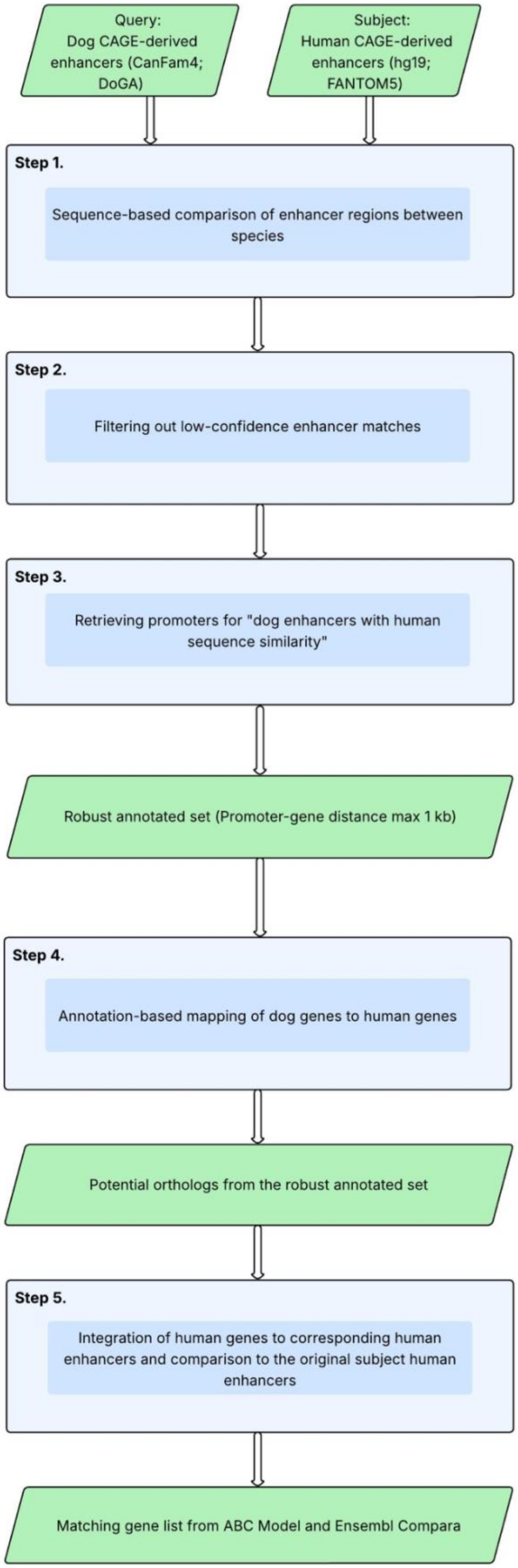

Figure S14: Workflow for Use Case 3: Enhancers exhibit sequence similarity between dogs and humans

The workflow illustrates the steps of the investigation of the enhancer sequence similarity between dogs and humans. Dog CAGE-derived enhancers (CanFam4; DoGA, *n* = 9,787) were compared to human CAGE-derived enhancers (hg19; FANTOM5, *n* = 65,423) using sequence similarity. Enhancer sequences were aligned with BLAST (dc-megablast), keeping the best human match per query, followed by filtering for low-confidence alignments (e-value $\leq 1\times10^{-3}$). Dog enhancers with human sequence similarity were intersected with an enhancer–promoter interaction dataset to identify associated promoters. Promoters were assigned using a robust annotation (<1 kb from gene). Genes linked to the identified promoters were mapped from dog to human using Ensembl Compara, retaining only one-to-one orthologs. This analysis identified 93 potential orthologs. From the original set of human enhancers, 1,199 with BLAST matches were retained and intersected with BEDtools and Activity-by-Contact (ABC) model enhancer–gene predictions, identifying 8,092 associated human genes. These genes were compared with 93 human orthologs identified from dog genes using Ensembl Compara, of which 76 overlapped in at least one cell line. The right panel shows a simplified overview of the workflow, and the left panel provides methodological details.

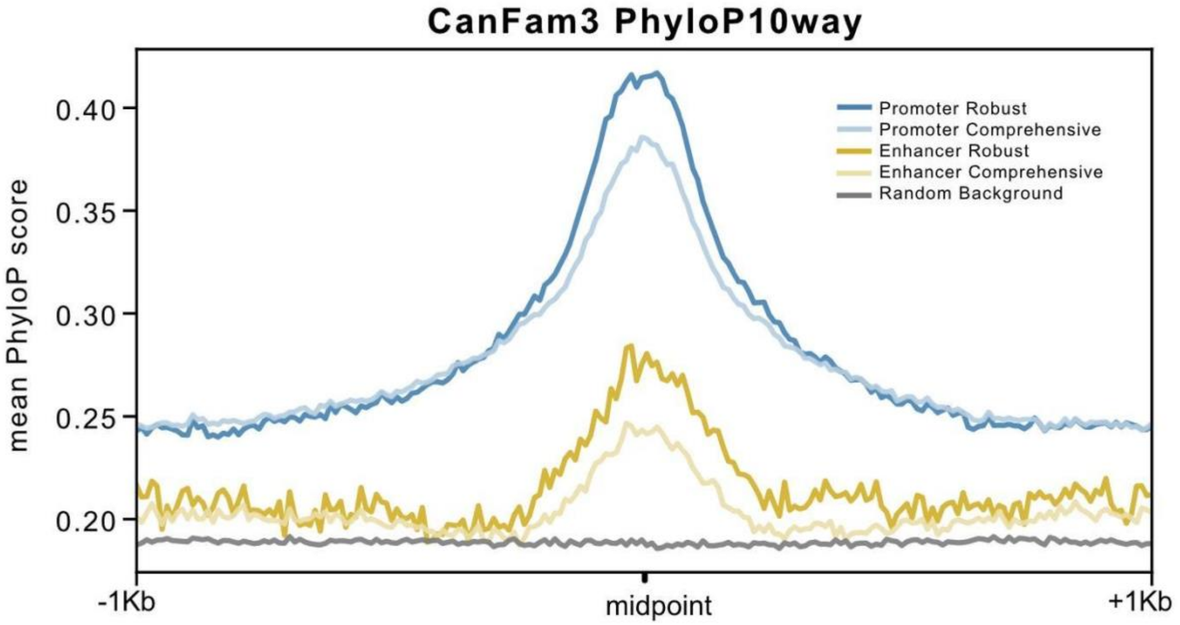


Figure S15 - Comparative Genomics: Promoter and Enhancer conservation

X-axes represent the mean PhyloP11 scores for dog genome regions derived from the study by Capriotti et al., which is based on alignments with 10 mammalian species genomes: Human,

Chimpanzee, Mouse, Rat, Cow, Panda, Marmoset, Cat, Horse, and Opossum. The conservation plot displays regions extending 1000 bp around the midpoints of each element in both directions.

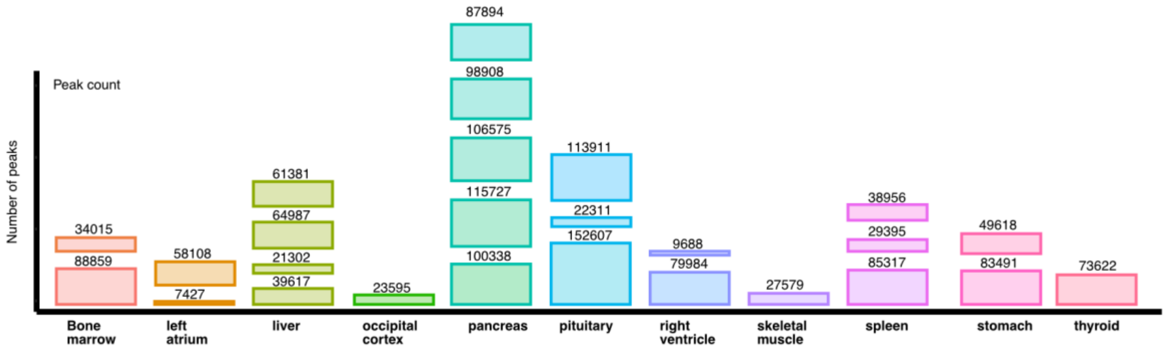


Figure S16 - ATAC-seq data processing Quality control

The X-axis shows the tissues provided by BarkBase. The y-axis represents the number of peaks. The occipital cortex was excluded from additional processing as a result of the low quality scores and the quantity of peaks.
